# 12h-clock control of central dogma information flow by XBP1s

**DOI:** 10.1101/559039

**Authors:** Yinghong Pan, Heather Ballance, Yisrael Schnytzer, Xi Chen, Oren Levy, Cristian Coarfa, Bokai Zhu

## Abstract

Our group recently discovered a cell-autonomous mammalian 12h-clock regulating physiological unfolded protein response. *Xbp1s* ablation impairs 12h-transcript oscillations *in vitro*, and we now show liver-specific deletion of *XBP1s* globally impaired murine 12h-transcriptome, but not the circadian rhythms *in vivo*. XBP1s-dependent 12h-transcriptome is enriched for transcription, mRNA processing, ribosome biogenesis, translation, and protein endoplasmic reticulum (ER)-Golgi processing/sorting in a temporal order consistent with the progressive molecular processing sequence described by the central dogma information flow (CEDIF). The 12h-rhythms of CEDIF are cell-autonomous and evolutionarily conserved in circatidal marine animals. Mechanistically, we found the motif stringency of promoter XBP1s binding sites, but not necessarily XBP1s expression, dictates its ability to drive 12h-rhythms of transcription and further identified GABP as putative novel transcriptional regulator of 12h-clock. We hypothesize the 12h-rhythms of CEDIF allows rush hours’ gene expression and processing, with the particular genes processed at each rush hour regulated by circadian and/or tissue specific pathways.

## INTRODUCTION

All life on earth is governed by biological rhythms that are defined as self-sustained oscillations cycling with a fixed period. Biological clocks enable organisms to keep track of the time of day and to adjust their physiology to recurring daily changes in the external environment, including nutrient and microenvironment status. Not surprisingly, lifestyle behaviors (e.g. shift work) that chronically de-regulate biological clocks are strongly associated with increased risk for a plethora of diseases, including but not limited to metabolic syndromes, cancer, sleep disorders, autoimmune diseases, brain disorders and cardiovascular diseases (Logan and McClung, 2019; Masri and Sassone-Corsi, 2018; Morris et al., 2016; Paganelli et al., 2018; Roenneberg and Merrow, 2016; Zhou et al., 2016).

Our understandings of biological rhythms in mammals have expanded beyond the well-characterized circadian rhythms (∼24h oscillation) in recent years through the discovery of the existence of 12h rhythms in mice (Hughes et al., 2009; Zhu et al., 2018). In a seminal 2009 study, using microarray to profile high resolution temporal mouse hepatic transcriptome, the authors originally identified approximately 200 genes cycling with a dominant 12h period, which are enriched in unfolded protein response (UPR) and endoplasmic reticulum (ER) stress pathway (Hughes et al., 2009). Since then, a handful of studies followed up on this initial discovery and proposed different hypothesis on the how the 12h rhythms are established (Cretenet et al., 2010; Hughes et al., 2012; Westermark and Herzel, 2013; Zhu et al., 2018). Early studies favor the hypothesis that the mammalian 12h rhythms are not cell-autonomous and instead are established by the combined effects of circadian clock and fasting-feeding cues. This conclusion was largely based upon the findings showing the lack of cell-autonomous 12h rhythms of gene expression in forskolin-synchronized NIH3T3 cells and altered 12h rhythms of gene expression under certain feeding and circadian clock perturbation conditions (Cretenet et al., 2010; Hughes et al., 2009; Hughes et al., 2012). Alternatively, it was suggested that two circadian transcription activators or repressors appearing in anti-phase is theoretically capable of establishing 12h rhythms of gene expression in a cell-autonomous manner (Westermark and Herzel, 2013). Contrary to these hypotheses, our group proposed a third scenario, whereby the mammalian 12h rhythms are not only cell-autonomous, but also are established by a dedicated ‘12h-clock’ separate from the circadian clock and function to coordinate cellular stress with metabolism (Antoulas et al., 2018; Zhu et al., 2018; Zhu et al., 2017).

Evidences supporting the existence of a cell-autonomous mammalian 12h-clock include *1)* the presence of intact hepatic 12h rhythms of ER and metabolism-related gene expression in *BMAL1* knock-out and *Clock*^Δ*19*^ mutant mice *in vivo* under free-running conditions (Zhu et al., 2018; Zhu et al., 2017); *2)* the detection of cell-autonomous 12h rhythms of gene expression in mouse embryonic fibroblasts (MEFs) (Zhu et al., 2018; Zhu et al., 2017); *3)* 12h rhythms of gene expression and metabolism can be established *in vitro* after synchronization by tunicamycin or glucose depletion in a *Bmal1*-independent manner (Zhu et al., 2018; Zhu et al., 2017); *4)* 12h rhythms of gene expression are highly conserved evolutionarily: they are found in species as divergent as crustacean, sea anemone, *C. elegans*, zebrafish and mammals including mouse and baboon (Zhu et al., 2018; Zhu et al., 2017) and *5)* 12h-cycling genes arose much earlier during evolution than circadian genes (averaged evolution age of 1,143 million years for 12h-cycling genes compared to 973 million years for circadian genes), implying a more ancient and distinct origin of 12h-clock from the circadian clock (Castellana et al., 2018).

Due to the strong enrichment of ER stress and UPR pathways in hepatic transcriptome exhibiting 12h rhythms, we initially hypothesized that the mammalian 12h-clock may be regulated transcriptionally by the UPR transcription factor XBP1s (Zhu et al., 2017). In agreement with this hypothesis, we previously found that siRNA-mediated knockdown of *Xbp1* in MEFs severely impaired cell-autonomous 12h oscillations of several genes including *Eif2ak3* and *Sec23b* (Zhu et al., 2018; Zhu et al., 2017). While these data suggest a role of XBP1s in regulating 12h rhythms of gene expression *in vitro*, whether XBP1s is a master transcriptional regulator of mammalian hepatic 12h-clock *in vivo* remains unknown. In the current study, we discovered that hepatic specific deletion of *XBP1s* severely impaired global hepatic 12h-transcriptome, without affecting the circadian clock nor fasting-feeding behavior in mice. XBP1s-dependent 12h-transcriptome is enriched for transcription, mRNA processing, ribosome biogenesis, translation, and protein processing/sorting in the ER-Golgi in a temporal order consistent with the progressive molecular processing sequence described by the central dogma information flow (CEDIF) and preferentially peaks during dawn and dusk in a day. In addition, 12h rhythms of CEDIF is coupled to 12h-rhythms of nucleotide and amino/nucleotide sugar metabolism. Importantly, the 12h rhythms of CEDIF is cell-autonomous and observed in serum-synchronized liver MMH-D3 cells *in vitro;* and evolutionarily conserved in marine animals possessing dominant circatidal clock. Mechanistically, we found the motif stringency of XBP1s binding sites, but not necessarily its expression, dictates its ability to drive 12h-rhythms of mRNA transcription. We further identified GA-binding protein (GABP) as a potential novel transcriptional regulator of mammalian 12h-clock acting downstream or in parallel with XBP1s. Based upon this evidence, we propose a vehicle-cargo hypothesis denoting the distinct functions of 12h versus circadian clock: whereas the 12h clock allows for elevated rush hours’ gene expression and processing by controlling the global traffic capacity of the central dogma information flow (thus the vehicle), the circadian clock and/or other tissue specific pathways dictate the particular genes processed at each rush hour (thus the cargo).

## RESULTS

### Liver-specific deletion of XBP1 does not alter rhythmic locomotor activity nor fasting-feeding cycles in mice

To delete XBP1 specifically in the liver, we crossed XBP1^Flox^ mice (Xbp1^*fl/fl*^ mice with loxP sites flanking exon 2 of the *Xbp1* gene) (Lee et al., 2008) with Albumin-CRE transgenic mice as previously described (Olivares and Henkel, 2015). Liver-specific deletion of XBP1s was confirmed by both qPCR and western blot analysis (Figures 1A-D). Consistent with previous reports (Cretenet et al., 2010; Zhu et al., 2017), robust 12h rhythms of total *Xbp1* as well as spliced form of *Xbp1* (*Xbp1s*) mRNA level were observed in XBP1^*Flox*^, but not in XBP1 liver specific knockout (XBP1^*LKO*^) mice, indicating XBP1s autoregulates its own 12h rhythm of expression (Figures 1A-C). We further observed a ∼3h phase delay between the acrophases of *Xbp1s* (0.3h) and total *Xbp1* (3.4h) (Figure 1C).

**FIGURE 1.**
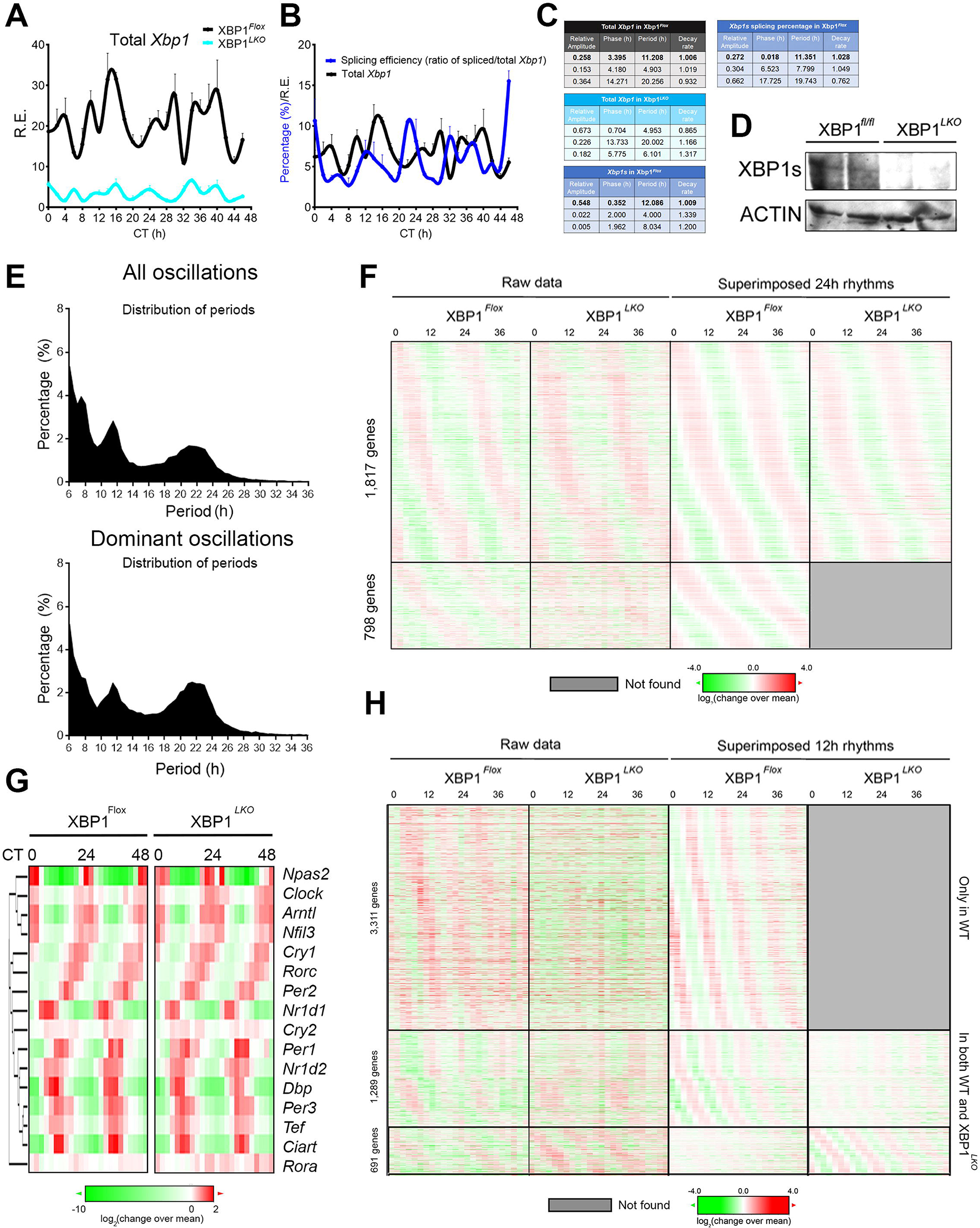
Liver-specific deletion of XBP1s impairs global hepatic 12h-transcriptome, but not the circadian rhythm in mice. **(A)** qPCR analysis of total hepatic *Xbp1* mRNA in wild-type (XBP1^*Flox*^) and XBP1 liver-specific knockout (XBP1^*LKO*^) mice. **(B)** Calculated splicing efficiency (ratio of *Xbp1s* to total *Xbp1* mRNA) and total hepatic *Xbp1* mRNA in XBP1^*Flox*^ mice. **(C)** Eigenvalue/pencil analysis of total *Xbp1* mRNA level in XBP1^*Flox*^ and XBP1^*LKO*^ mice, *Xbp1s* mRNA level and the splicing efficiency in XBP1^*Flox*^ mice. **(D)** Western blot analysis of total hepatic XBP1s in XBP1^*Flox*^ and XBP1^*LKO*^ mice. **(E)** Distribution of periods of all (top) and dominant oscillations (bottom) identified by eigenvalue/pencil method from XBP1^*Flox*^ mice. **(F)** Heat map of all circadian gene expression (or lack thereof) in XBP1^*Flox*^ and XBP1^*LKO*^ mice with both raw data and superimposed 24h rhythms shown. **(G)** Heat map of 16 core circadian clock gene expression in XBP1^*Flox*^ and XBP1^*LKO*^ mice with raw data shown. **(H)** Heat map of all ∼12h gene expression (or lack thereof) in XBP1^*Flox*^ and XBP1^*LKO*^ mice with both raw data and superimposed ∼12h rhythms shown. Data are graphed as the mean ± SEM (n = 3 to 5).

To rule out the potential effects of liver specific ablation of XBP1s on mouse locomotor activity and/or feeding behavior, which could confound the interpretation of the transcriptome data, we subjected both XBP1^Flox^ and XBP1^*LKO*^ mice to home cage and CLAMS, respectively. As shown in Figures S1A-E, liver-specific deletion of XBP1 does not alter rhythmic locomotor activity nor fasting-feeding cycles in mice.

### Liver-specific deletion of XBP1 does not affect the core circadian clock in mice

To identify XBP1s-dependent oscillating transcriptome, we performed RNA-Seq analysis in the liver of XBP1^*Flox*^ and XBP1^*LKO*^ mice at 2h interval for a total of 48 hours under constant darkness in duplicates (Table S1 and see Materials and Methods for details). All superimposed oscillations in either XBP1^Flox^ or XBP1^*LKO*^ mice were identified by the newly developed eigenvalue/pencil method (Antoulas et al., 2018; Zhu et al., 2017), which unlike canonical oscillation-identification methods such as JTK_CYCLE and ARSER, does not require a pre-assignment of period range and thus allow unbiased identification of all superimposed oscillations (Antoulas et al., 2018; Zhu et al., 2017). Consistent with past findings (Zhu et al., 2017), the vast majority of oscillations identified were circadian rhythms and oscillations that cycle at the second (∼12h) and third harmonics (∼8h) of the circadian rhythm [due to the 2h sampling frequency of the current study, only up to third harmonics can be identified with high confidence (Antoulas et al., 2018)] (Figure 1E and Table S2). To determine the false discovery rate (FDR) of the identified rhythmic transcripts, we used a permutation-based method that randomly shuffles the time label of gene expression data and subjects each of the permutation dataset to eigenvalue/pencil method as previously described (Rey et al., 2018). As expected, permutation datasets are devoid of distinct populations of oscillations cycling at different harmonics of the circadian rhythm (Figures S2A-B). In this way, we identified a total of 2,615 ∼24h circadian genes with FDR of 0.05, 5,299 ∼12h-cycling genes with FDR of 0.24 and 5,608 ∼8h-cycling genes with FDR of 0.31 respectively (Table S3, S4 and see Materials and Methods for details) in XBP1^*Flox*^ mice. Agreeing with past findings (Zhu et al., 2017), while the phases of circadian rhythms are evenly distributed throughout the day (Figure S3B), the phases of ∼12h rhythms are enriched at dawn (CT0-2) and dusk (CT12-14), which is more evident for dominant oscillations (Figures S3H).

Of all circadian genes identified in XBP1^*Flox*^ mice, ∼70% were unaffected by hepatic XBP1s ablation, which includes all known core circadian clock genes such as *Arntl1* (*Bmal1*), *Clock, Npas2, Per1, Per2, Cry1*, Cry2, *Nr1d1, Nr1d2, Nfil3, Dbp* and *Tef* (Figures 1F,G S2C). On average, comparable average gene expression and phase distribution of circadian oscillations are found between XBP1^*Flox*^ and XBP1^*LKO*^ mice (Figures S3A-C). Circadian genes not affected by hepatic XBP1s ablation are enriched in circadian rhythm and metabolic pathways (Figures S3D, E), which are the dominant biological pathways under hepatic circadian clock control (Eckel-Mahan and Sassone-Corsi, 2013; Stashi et al., 2014). We further identified 798 genes, whose superimposed circadian rhythms are abolished in XBP1^*LKO*^ mice (Figure 1F). These genes are enriched in biological pathways including RNA degradation and PI3K-AKT signaling, processes not known be under strong circadian clock control (Figure S3F, G). Further, circadian rhythms of these genes oscillate with a much smaller amplitude and are often subordinate oscillations (Figures 1F, S3G). We conjecture the circadian rhythms of these genes are likely not driven by the circadian clock and are altered as an indirect consequence of XBP1s ablation. Taken together, our data indicate that the liver-specific deletion of XBP1 does not affect the core circadian clock in mice.

### Liver-specific deletion of XBP1 globally impairs 12h transcriptome, which is enriched in pathways regulating central dogma information flow

In sharp contrast to largely intact circadian rhythms in XBP1^*LKO*^ mice, ablation of XBP1 in the liver significantly impairs the global 12h transcriptome profile (Figures 1H, S2D). Specifically, we identified 3,311 genes (62.5%), 1,289 genes (24.4%) and 691 genes (13.1%), whose superimposed 12h rhythms were abolished, dampened or increased in the absence of XBP1, respectively (Figure 1H). Subsequent Gene ontology (GO) analysis on those 4,600 (86.9%) genes whose 12h rhythms were either abolished or dampened in XBP1^*LKO*^ mice revealed top enriched KEGG pathways of protein processing in the ER, RNA transport, splicesome, mRNA surveillance, protein export and ribosome biogenesis (Figures 2A, B, Table S5). In addition, we also found 12h genes enriched in Golgi apparatus by using a different GO category (Figure 2B, Table S5). These enriched biological pathways are reminiscent of the progressive molecular processing sequence described by the central dogma information flow (CEDIF), namely, mRNA processing in the nucleus, ribosome biogenesis/translation in the nucleus/cytosol, protein processing and sorting in the ER/Golgi apparatus in a temporal order (Figure 2C). Importantly, we found both anabolism and catabolism pathways are enriched, suggesting an overall XBP1s-dependent 12h-rhythm of RNA and protein processing rate.

**FIGURE 2.**
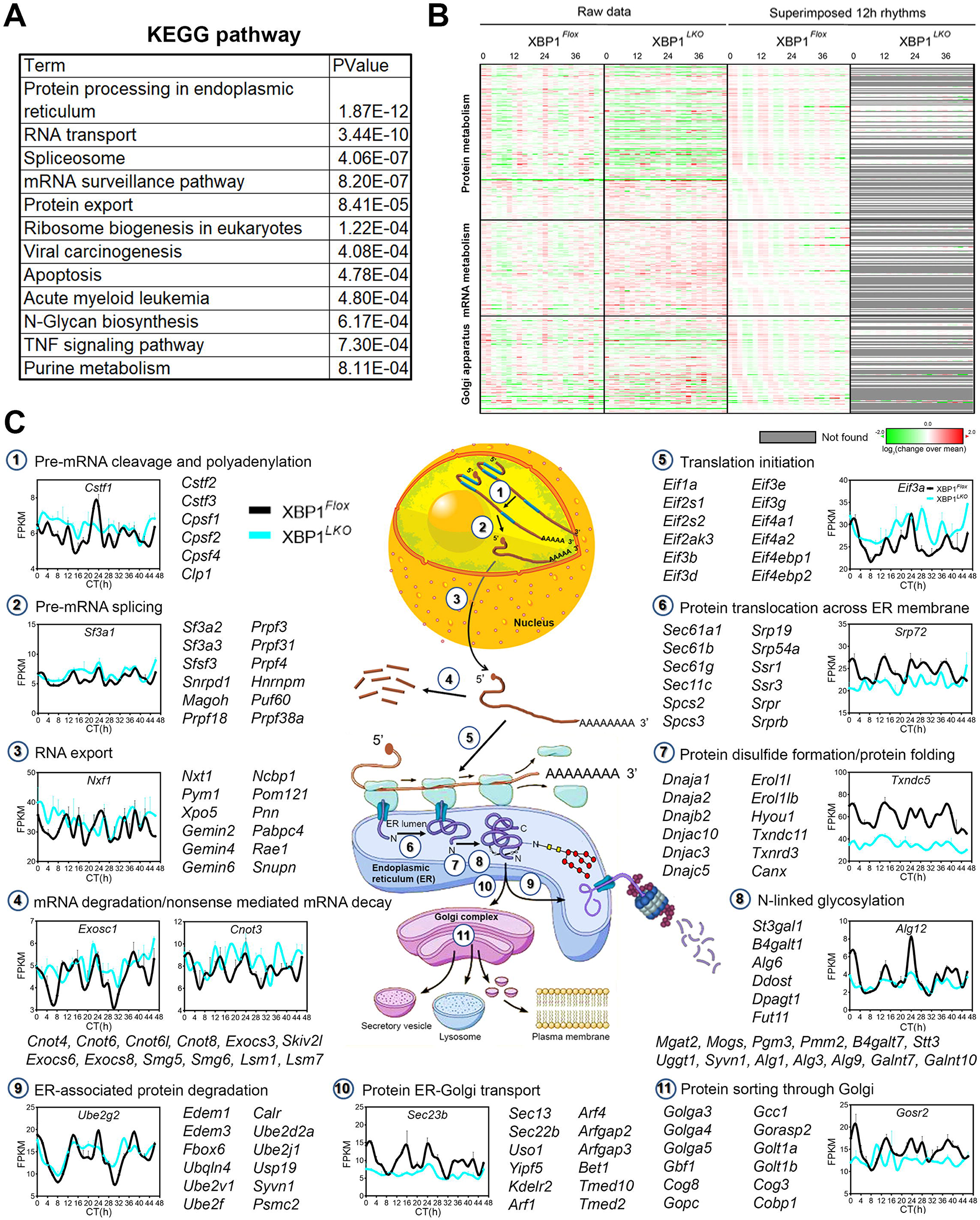
XBP1s-dependent hepatic 12h transcriptome is enriched in regulating central dogma information flow. **(A)** GO analysis showing enriched KEGG pathways and their corresponding p values for XBP1s-dependent 12h transcriptome. **(B)** Heat map of ∼12h-cycling gene expression (or lack thereof) involved in mRNA metabolism, protein metabolism and the Golgi apparatus in XBP1^*Flox*^ and XBP1^*LKO*^ mice with both raw data and superimposed ∼12h rhythms shown. **(C)** Diagram illustrating each steps involved in the central dogma information flow from mRNA processing all the way to protein sorting in the Golgi, and RNA-Seq data for representative genes in XBP1^*Flox*^ and XBP1^*LKO*^ mice. Additional selected gene names belonging to each step are also shown. Data are graphed as the mean ± SEM (n = 2).

Based upon the GO analysis, we divided the XBP1s-dependent 12h-cycling CEDIF into eleven steps and carefully annotated the genes and assigned them into one of the eleven steps (Figure 2C). In total, we identified 114 genes involved in RNA metabolism (Figure 2C and Table S6). These genes encompass the steps of pre-mRNA cleavage and polyadenylation, pre-mRNA splicing, RNA export and mRNA degradation and nonsense-mediated mRNA decay, with prominent examples including cleavage stimulatory factor (*Cstf*) and cleavage and polyadenylation specificity factor (*Cpsf*) family members involved in pre-mRNA cleavage and polyadenylation (Mandel et al., 2008), numerous mRNA splicing factors including *Sf3a1-3* and *Prpf* family members, small nuclear ribonucleoproteins that are part of splicesome (Maniatis and Reed, 1987) (*Snrpa, Snrpd1, Snrpe*), RNA export factors *Nxf1* and *Nxt1*, multipotent gene expression regulator CCR4-NOT complex family member (Collart, 2016) (*Cnot3, Cnot4, Cnot6*) and exosome complex components mediating RNA degradation (Zinder and Lima, 2017) (*Exosc1, Exosc3*) (Figure 2C). Overall, we found the mean expression of these 12h cycling RNA metabolism genes were elevated in XBP1^*LKO*^ mice (Figure 2B).

The most dominant GO pathways whose 12h rhythms are dependent on XBP1s are protein metabolism, which include a total of 170 genes (Figure 2C and Table S6). These pathways include ER-associated protein degradation (*Edem1* and *Edem3*), translation initiation (eukaryotic initiation factor members), protein translocation across the ER membrane (the ER translocon Sec61 proteins), protein folding in the ER (HSP40 family members), protein glycosylation in the ER (multiple ALG members) and protein transport from ER to the Golgi (COPII subunits *Sec13, Sec22* and *Sec23*). For these genes, the average expression was reduced in XBP1^*LKO*^ mice (Figure 2B).

The last category includes 110 genes involved in maintaining the integrity of the structure and function of the Golgi apparatus, where additional protein and lipid transport and protein N-linked glycosylation also take place (Figure 2C and Table S6). Examples include golgins family members (*Golga3-5*) playing key roles in the stacking of Golgi cisternae (Barr and Short, 2003), and Golgi transport protein (*Golt1a and Golt1b*) involved in fusion of ER-derived transport vesicles with the Golgi complex (Conchon et al., 1999). For these genes, the average expression was similar between XBP1^*Flox*^ and XBP1^*LKO*^ mice (Figure 2B).

To determine whether nuance exists in the 12h transcriptome profile of genes involved in different steps of CEDIF, we performed t-distributed Stochastic Neighbor Embedding (t-SNE) analysis on the superimposed 12h transcriptome of XBP1^*Flox*^ mice revealed by the eigenvalue/pencil method (Figure 3A). t-SNE has been extensively used in the analysis of single-cell RNA-Seq data to reveal local structure in high-dimensional data and to cluster different single cells based upon their gene signature through dimension reduction (Yang et al., 2018). Here we reason that if such nuance exists, these subtle differences must be encoded by the shape of a waveform, specifically, by the different combinations of amplitude, phase, decay rate, and period information of the superimposed 12h rhythms; and therefore, are likely to be revealed through the spatial clustering of genes belonging to different steps of CEDIF based upon their 12h transcriptome profile. As expected, t-SNE analysis revealed that the clustering of CEDIF genes exhibits a spatial trajectory consistent with the direction of central dogma information flow (Figures 3B-C, S4A-C). This is reflected both by a gradual spatial separation of genes involved in mRNA processing, ribosome biogenesis/translation, and protein processing in the ER/Golgi (Figures 3B, S4A-B), and by a much finer spatial separation of genes involved in the different sub steps of protein metabolism (Figures 3C, S4C). Lastly, we found one contributing factor to the spatial gene clustering is the progressive phase delay of genes involved in mRNA processing, protein processing in the ER, and protein sorting in the Golgi, changing from CT0 to CT3 and from CT12 to CT15. This progressive phase delay is consistent with the direction of central dogma information flow (Figure 3D). In sum, these data indicate a delicately-orchestrated 12h rhythms of CEDIF by XBP1s.

**FIGURE 3.**
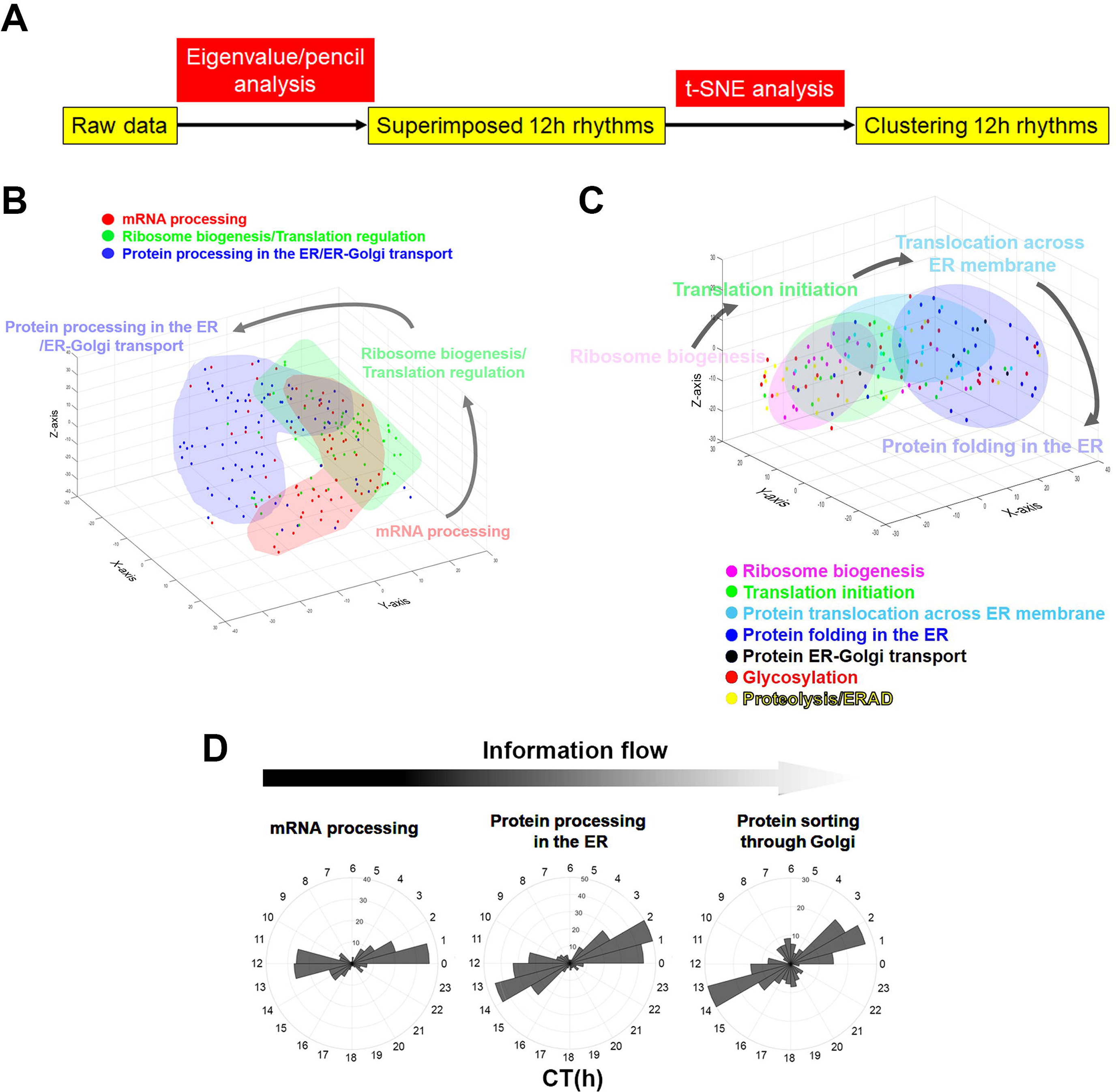
Clustering CEDIF genes based upon their 12h transcriptome profile reveals a spatial order consistent with the direction of central dogma information flow. **(A)** Flow chart illustrating the framework for clustering CEDIF genes based upon their superimposed 12h transcriptome profile. **(B)** Clustering of genes involved in mRNA processing, ribosome biogenesis/translation initiation and protein processing and transport based upon their superimposed 12h transcriptome projected onto 3D t-SNE space. **(C)** Clustering of genes involved in different sub-steps of protein metabolism based upon their superimposed 12h transcriptome projected onto 3D t-SNE space. **(D)** Polar histograms demonstrating phase distributions of genes involved in different steps of central dogma information flow.

### Coordinated 12h rhythms of nucleotide and nucleotide/amino sugar metabolism and CEDIF

In addition to biological pathways involved in CEDIF, we also observed statistically significant enrichment of multiple metabolic pathways in XBP1s-dependent 12h genes (Figure 4A). Two notable pathways are nucleotide metabolism (purine and pyrimidine metabolism), and amino sugar and nucleotide sugar/N-glycan biosynthesis, which provide metabolite precursors for protein N-linked glycosylation in the ER and Golgi (Figures 4A-C). In particular, nine of the eleven genes encoding metabolic enzymes for *de novo* purine synthesis, and five of the six genes encoding enzymes for *de novo* pyrimidine synthesis exhibit XBP1s-dependent 12h oscillation, which include rate-limiting enzymes PPAT and CAD, for purine and pyrimidine biosynthesis, respectively (Figure 4C). In addition, several genes encoding enzymes for purine salvage pathways including XDH and GDA also reveal XBP1s-dependent 12h oscillation (Figure 4C). Intriguingly, we also uncovered twelve genes encoding subunits of RNA polymerase I, II and III exhibiting XBP1s-dependent 12h rhythms (Figure 4C). To determine whether 12h rhythms of hepatic metabolic enzyme expression are associated with 12h hepatic metabolite oscillations, we performed *post hoc* analysis of a previously published high resolution (at 1h interval for a total of 48 hours) murine hepatic metabolome dataset using the eigenvalue/pencil method (Krishnaiah et al., 2017), and found 12h oscillations of various nucleoside and nucleotide and the N-linked glycosylation precursor UDP-N-Acetyl-Amino sugar (Figure 4D and Table S7).

**FIGURE 4.**
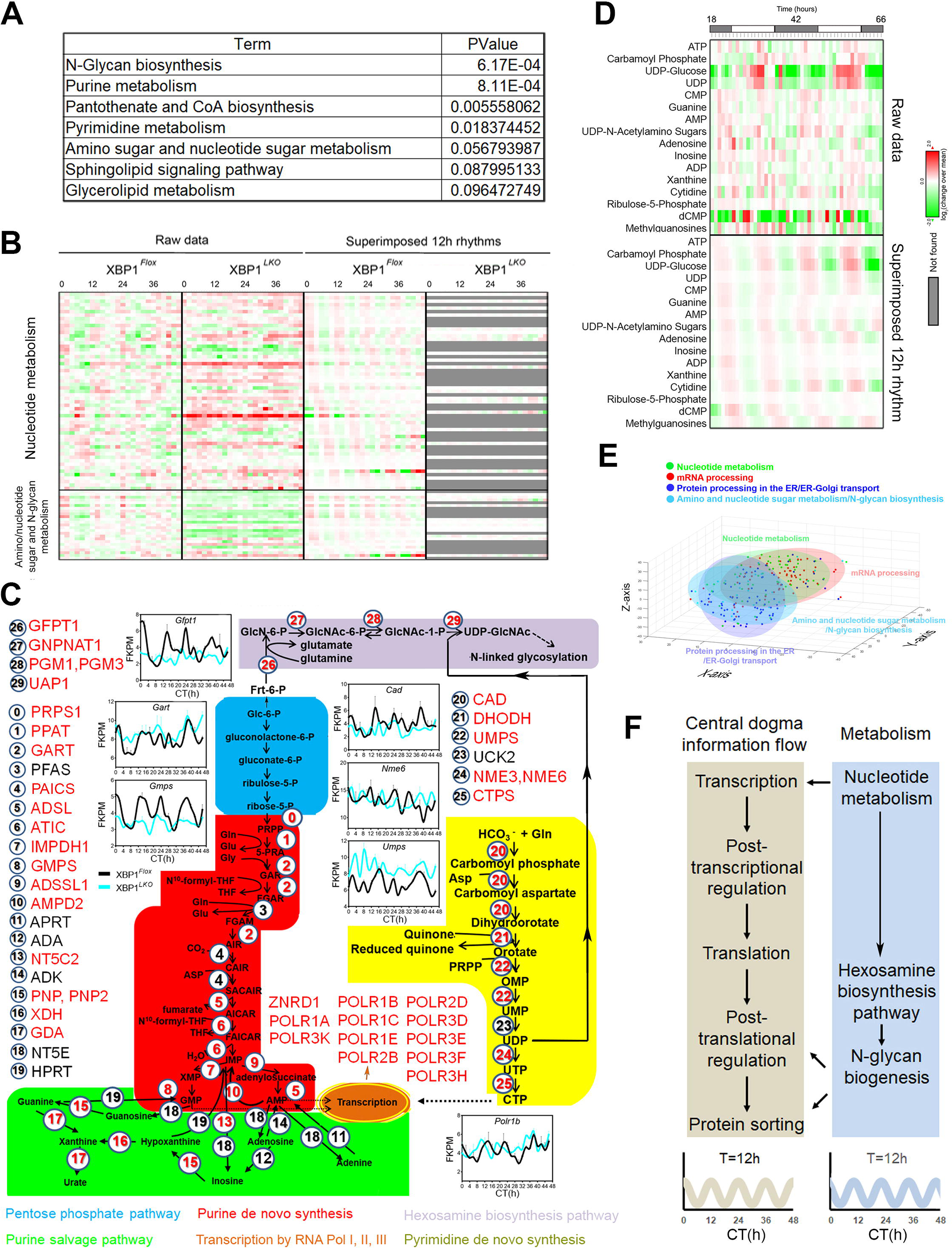
Coordinated 12h rhythms of nucleotide and nucleotide/amino sugar metabolism with 12h rhythms of CEDIF. **(A)** GO analysis showing enriched KEGG metabolic pathways and their corresponding p values for XBP1s-dependent 12h transcriptome. **(B)** Heat map of ∼12h-cycling gene expression (or lack thereof) involved in nucleotide, amino/nucleotide sugar and N-Glycan metabolism in XBP1^*Flox*^ and XBP1^*LKO*^ mice with both raw data and superimposed ∼12h rhythms shown. **(C)** The diagram illustrates key metabolic pathways (highlighted by different colors) and corresponding genes that are related to nucleotide and amino/nucleotide sugar metabolism. Genes are numbered from 0 to 29 and those highlighted in red exhibit XBP1s-dependent 12h oscillation. RNA-Seq data for representative genes are shown. **(D)** Heat map of mouse 12h cycling hepatic metabolites related to nucleotide and amino/nucleotide sugar metabolism identified by the eigenvalue/pencil method from published metabolomic dataset (Krishnaiah et al., 2017). Both the raw data as well as 12h superimposed oscillations are shown. **(E)** Clustering of genes involved in different metabolic pathways and CEDIF based upon their superimposed 12h transcriptome projected onto 3D t-SNE space. **(F)** A model demonstrating coordinated 12h rhythms of nucleotide and nucleotide/amino sugar metabolism with 12h rhythms of CEDIF Data are graphed as the mean ± SEM (n = 2).

We hypothesize that the 12h rhythmic expression of genes regulating nucleotide and amino sugar/nucleotide sugar metabolism are commensurate with those regulating mRNA and protein processing, respectively (Figure 4F). To test this hypothesis, we again performed t-SNE analysis on the superimposed 12h transcriptome of these four pathways. As shown in Figures 4E and S4D, we observed that genes involved in nucleotide metabolism and mRNA processing occupy largely overlapping t-SNE space, while genes involved in amino sugar/nucleotide sugar/N-glycan biosynthesis and protein processing in the ER/Golgi also reveal similar spatial clustering. Together, we discovered a coordinated 12h rhythms of nucleotide and nucleotide/amino sugar metabolism and CEDIF in mouse liver (Figure 4F).

### Cell-autonomous 12h rhythms of CEDIF and metabolism gene expression in hepatocyte and MEF

We wondered whether the 12h rhythms of CEDIF and metabolism gene expression observed *in vivo* are also cell-autonomous, namely, whether they can also be observed in cell culture systems deprived of systemic cues. We addressed this question first by performing a *post hoc* analysis of time series transcriptome of serum-synchronized murine liver MMH-D3 cell (Atwood et al., 2011). Upon examining the raw data, we observed noticeable baseline changes in most reported mRNA oscillations. Therefore, we subjected the raw data to polynomial detrend before the identification of all superimposed oscillations using the eigenvalue/pencil analysis (See Materials and Methods for details). Similar to what is observed *in vivo*, the majority of oscillations identified in MMH-D3 cells were circadian rhythms (which cycle at a slightly shorter period of 21.6h) and oscillations that cycle at the second (∼10.8h) and third harmonics (∼7.3h) of the circadian rhythm, although more oscillations cycling with periods in between were observed *in vitro* compared to *in vivo* (Figures S5A, B, Table S8). Genes with circadian oscillations were enriched in biological pathways of circadian rhythm and HIF-1 signaling and include core circadian clock genes such as *Bmal1, Clock, Nr1d1, Nr1d2* and *Per1* (Figures S5C-E, Table S8), consist with past findings (Adamovich et al., 2017; Atwood et al., 2011; Peek et al., 2017; Wu et al., 2017).

Enriched biological pathways associated with ∼12h genes (9.5h to 12.5h) in MMH-D3 cells reveal large convergence with those found in mouse liver *in vivo*, including all CEDIF pathways: mRNA processing (splicesome and mRNA surveillance pathways), translation and protein metabolism (ribosome, ubiquitin-mediated proteolysis, protein processing in the ER, SNARE interactions in vesicular transport, protein export) and Golgi apparatus (Figures 5A,B and Table S9). In total, 228 12h-cycling CEDIF genes were commonly found in mouse liver *in vivo* and MMH-D3 cells *in vitro* (Figures 5C, D and Table S10) and representative genes were shown in Figures 5G and S5G, which also include the 12h oscillation of total *Xbp1* mRNA expression. After converting the time post serum shock into CT time, we further observed a similar bimodal phase distribution of those 12h-cycling CEDIF genes at dawn and dusk (Figure 5E). For the other 273 12h-cycling CEDIF genes only found in MMH-D3 cells *in vitro*, they often represent different members of the same gene family as found *in vivo* (Figures 5C and S5H). t-SNE analysis on the 12h transcriptome of MMH-D3 CEDIF genes reveal spatial separation of mRNA processing genes with those regulating protein processing (Figures 5F and S5F), in line with *in vivo* findings.

**FIGURE 5.**
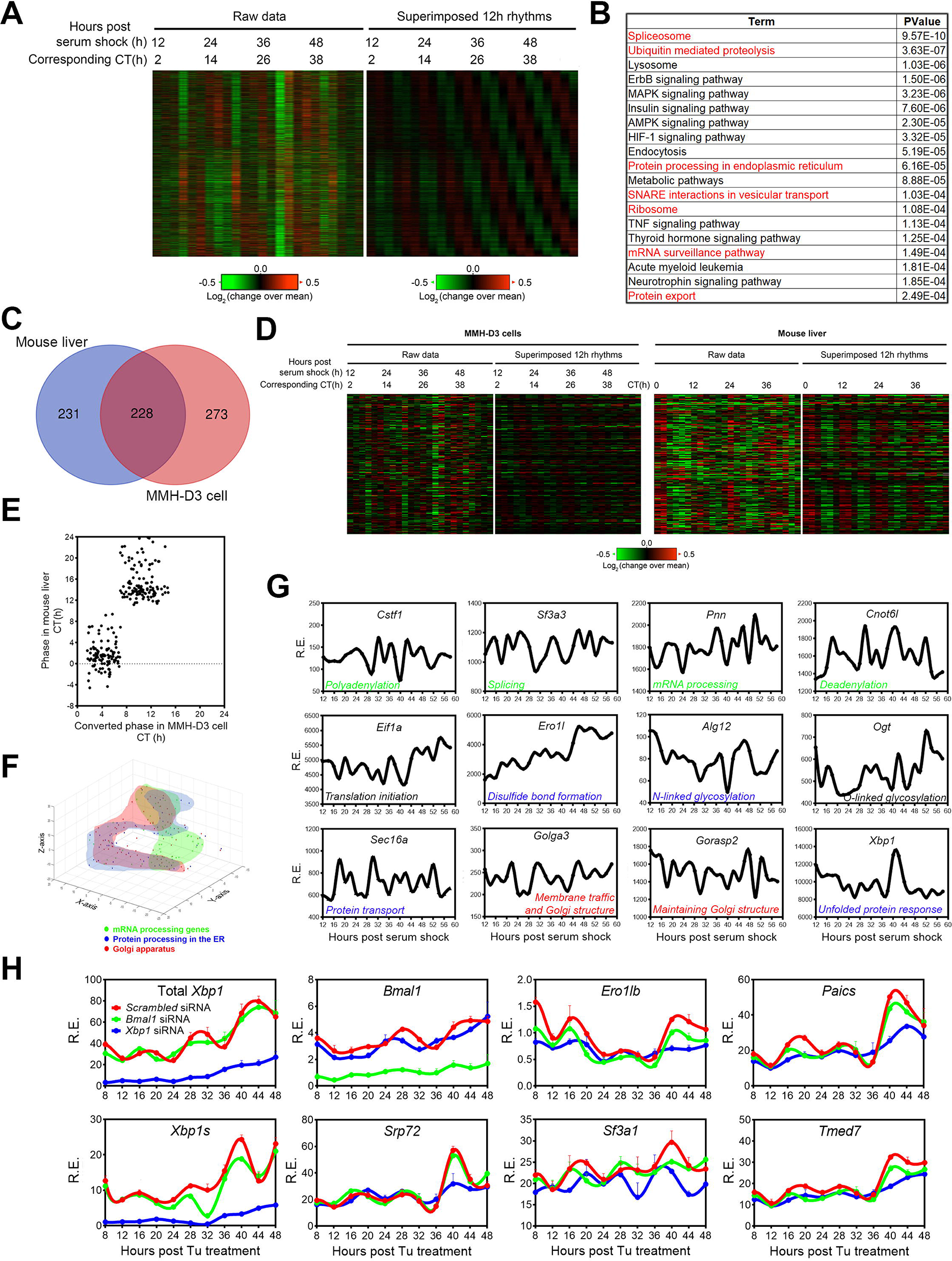
The 12h-rhythms of CEDIF gene expression are cell-autonomous. **(A)** Heat map of all ∼12h-cycling gene expression uncovered from serum shock-synchronized murine liver cell line MMH-D3 identified by eigenvalue/pencil analysis (Atwood et al., 2011), with both raw data and superimposed ∼12h rhythms shown. Both the original time after serum shock, as well as converted time in CT are shown. **(B)** GO analysis showing enriched KEGG pathways and their corresponding p values for cell-autonomous 12h transcriptome, with GO terms related to CEDIF highlighted in red. **(C)** Venn diagram comparison of 12h transcriptome involved in CEDIF from mouse liver *in vivo* and MMH-D3 cells *in vitro*. **(D)** Heat map of side-by-side comparison of CEDIF-related 12h gene expression in MMH-D3 cells and mouse liver, with both raw data and superimposed ∼12h rhythms shown. **(E)** Scatter plot comparing phases of CEDIF-related 12h gene oscillation in MMH-D3 cells and mouse liver. The phases of 12h oscillations in MMH-D3 are converted to corresponding time in CT. **(F)** Clustering of MMH-D3 genes involved in different CEDIF based upon their superimposed 12h transcriptome projected onto 3D t-SNE space. **(G)** Microarray data of representative ∼12h-cycling genes involved in CEDIF in MMH-D3 cells compiled from (Atwood et al., 2011). **(H)** MEFs were transfected with different siRNAs and treated with Tu (25ng/ml) for 2h and qPCR was performed at different time points post-Tu shock. Data are graphed as the mean ± SEM (n = 3).

To determine whether XBP1s also transcriptionally regulates 12h rhythms of CEDIF gene expression independently from the circadian clock *in vitro*, we knocked down *Xbp1* or *Bmal1* using siRNA and performed qPCR in dexamethasone or tunicamycin-synchronized MEFs as previously described (Zhu et al., 2017). In agreement with the unaffected core circadian clock in XBP1^*LKO*^ mice liver, knocking down *Xbp1* has no effects on dexamethasone-synchronized circadian oscillations of *Per2* and *Reverb*α (*Nr1d1*) expression, whose rhythms are nonetheless significantly impaired by *Bmal1* knock down as expected (Figure S5I). In tunicamycin-synchronized MEFs, robust 12h rhythms of both total and spliced *Xbp1* expression were observed, with a similar ∼3.5h phase difference found between the two as *in vivo* (Figures 5H and 1C). We further observed *Xbp1s*-dependent 12h rhythmic expression of genes involved in different steps of CEDIF (*Sf3a1* in pre-mRNA splicing, *Srp72* in protein translocation across ER membrane, *Ero1lb* in protein folding in the ER, *Tmed7* in protein transport in the cis-Golgi network), as well purine *de novo* synthesis gene *Paics.* These 12h rhythms are not affected by *Bmal1* knock down (Figures 5H and S5J). These evidence strongly support the existence of cell-autonomous 12h rhythms of CEDIF and metabolism gene expression.

### GABP as putative novel transcription regulators of mammalian 12h rhythm

XBP1s is known to act as a basic region leucine zipper (bZIP) transcription factor activating gene expression by binding to gene regulatory regions harboring consensus DNA sequence GCCACGT under ER stress condition (Chen et al., 2014; He et al., 2010; Kanemoto et al., 2005). Since hepatic XBP1s expression exhibits a 12h rhythm under physiological conditions without exogenous ER stress at both the mRNA and protein level (Figures 1A-C) (Cretenet et al., 2010; Zhu et al., 2017), we performed hepatic XBP1s ChIP-seq (at a 4h interval for a total of 48 hours) to globally profile its cistrome under constant darkness condition and identified a total of 3,129 high confidence binding sites (See Materials and Methods for details) (Figure 6A and Table S11). Consistent with its oscillating expression, XBP1s cistrome cycles with a 12h period, with peak binding observed at CT0, CT12, CT24 and CT36 (Figures 6A, C, S6D, F).

**FIGURE 6.**
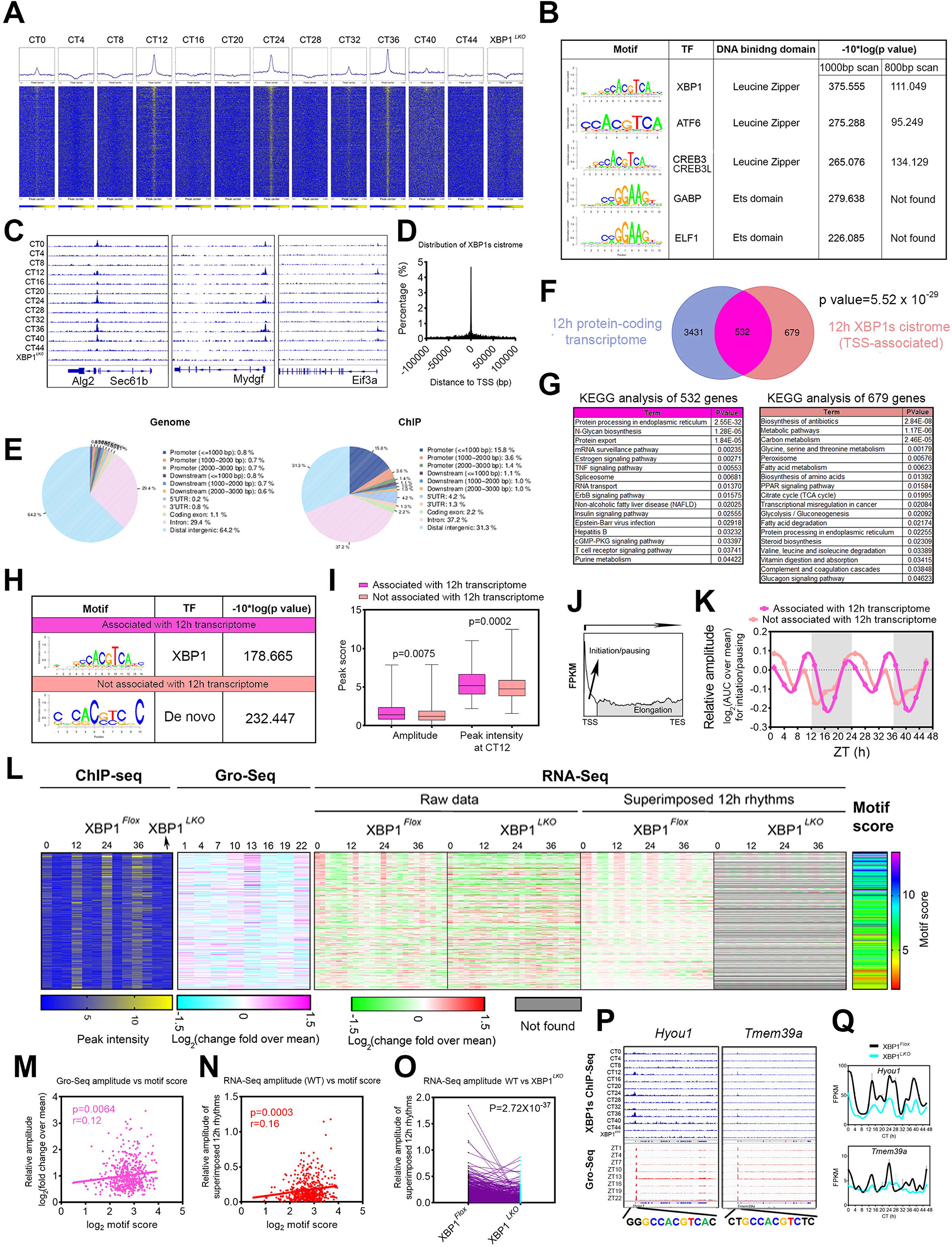
The motif stringency of XBP1s promoter binding sites dictates XBP1s’ ability to drive 12h-rhythms of transcription of CEDIF genes. **(A**) Heat maps of all XBP1s binding signal at 4h-interval in XBP1^*Flox*^ mice as well as in XBP1^*LKO*^ mice surrounding the center of XBP1s binding sites. **(B)** Top enriched SeqPos motifs common to 12h-cycling XBP1s cistrome. **(C)** Snapshot of target genes selected for alignment of XBP1s binding sites at different CTs in liver tissue of XBP1^*Flox*^ and XBP1^*LKO*^ mice. **(D)** Distribution of the distance of all XBP1s binding sites to transcription start site (TSS). **(E)** Distribution of mouse genome (left) and XBP1s cistrome (right) relative to known target genes. **(F)** Venn diagram depicting common and unique 12h-cycling protein-coding transcriptome and TSS-associated XBP1s cistrome. **(G)** GO analysis showing enriched KEGG pathways and their corresponding p values for 532 genes with both 12h-cycling transcriptome and XBP1s cistrome (left) and 679 genes with XBP1s cistrome but no 12h-cycling transcriptome (right). **(H)** Top enriched SeqPos motifs common to XBP1s cistromes associated with or without 12h-cycling transcriptome. **(I)** Calculated amplitude of cycling XBP1s binding signal and peak intensity of XBP1s binding at CT12 for XBP1s cistrome associated with or without 12h-cycling transcriptome. **(J)** A representative diagram depicting a typical Gro-Seq signal from TSS to transcription termination site (TES) of a gene and using area under curve (AUC) to calculate both transcription initiation/pausing and transcription elongation rates. **(K)** Log_2_ mean normalized transcription initiation rates calculated from the Gro-Seq data (Fang et al., 2014) for XBP1s target genes with or without associated 12h transcriptome. The data is double-plotted for better visualization. **(L-Q)** 532 genes with both proximal promoter XBP1s binding and 12h transcriptome in XBP1^*Flox*^ mice. **(L)** Comparisons of heat maps of XBP1s binding intensity, transcription initiation rates calculated from Gro-Seq (Fang et al., 2014), 12h-cycling gene expression (or lack thereof) in XBP1^*Flox*^ and XBP1^*LKO*^ mice with both raw data and superimposed ∼12h rhythms shown, and XBP1s binding motif score. **(M)** Scatter plot of Log_2_ transform of XBP1s binding motif score versus the relative amplitude of mRNA transcription initiation rates calculated from the Gro-Seq (Fang et al., 2014) for each gene in XBP1^*Flox*^ mice is shown, together with correlation coefficient *r* and p value that *r* is significantly different than 0. **(N)** Scatter plot of Log_2_ transform of XBP1s binding motif score versus the relative amplitude of superimposed 12h rhythms for each gene in XBP1^*Flox*^ mice is shown, together with correlation coefficient *r* and p value that *r* is significantly different than 0. **(O)** Relative amplitude of superimposed 12h rhythms for each gene in XBP1^*Flox*^ and XBP1^*LKO*^ mice. If a superimposed 12h rhythm is not found in XBP1^*LKO*^ mice, then an amplitude of 0 is used. **(P)** Snapshot of target genes selected for alignment of XBP1s binding sites at different CTs in liver tissue of XBP1^*Flox*^ and XBP1^*LKO*^ mice as well as published Gro-Seq data (Fang et al., 2014). Consensus XBP1s binding motifs located at each gene promoter are also shown. **(Q)** RNA-Seq data for representative genes in XBP1^*Flox*^ and XBP1^*LKO*^ mice. Data are graphed as the box and whisker plot (min to max) in **I** and mean ± SEM in **Q**.

**FIGURE 7.**
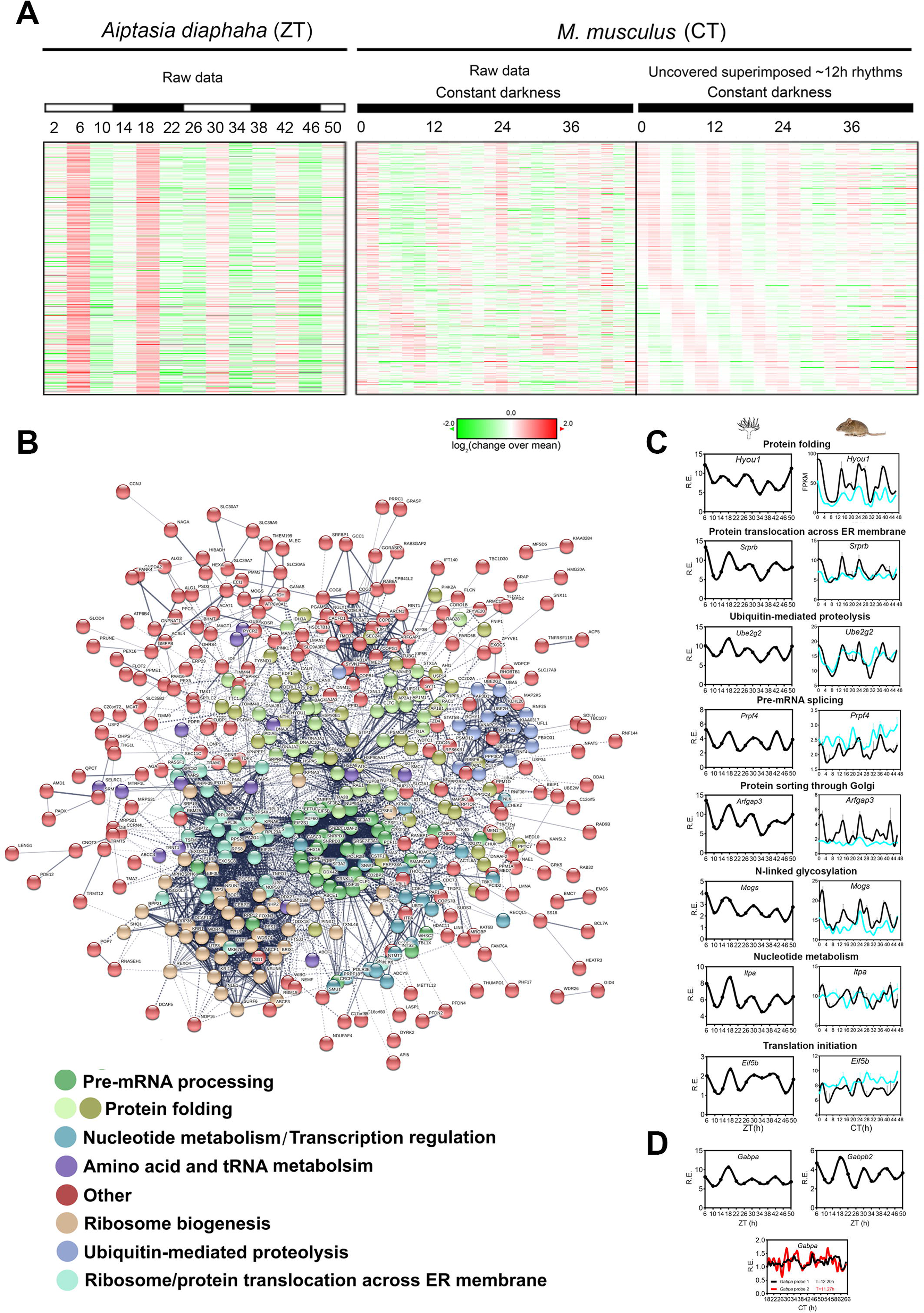
The 12h-rhythms of CEDIF gene expression are evolutionarily conserved in the aposymbiotic sea anemone *A. diaphaha*, which possesses a dominant circatidal clock. **(A)** Heat map of side-by-side comparison of evolutionarily conserved 12h gene expression in aposymbiotic *A. diaphaha* (Sorek et al., 2018) and mouse liver, with both raw data and superimposed ∼12h rhythms shown for the mouse liver data. **(B)** Predicted interactive network construction of these conserved 12h-cycling genes using STRING (Szklarczyk et al., 2015). Genes involved in different biological pathways are colored differently. **(C)** RNA-Seq data for representative genes in *A. diaphaha* (Sorek et al., 2018) and XBP1^*Flox*^ and XBP1^*LKO*^ mice. Data are graphed as the mean ± SEM (n = 2). **(D)** *Gabpa* and *Gabpb2* expression in aposymbiotic *A. diaphaha* (Sorek et al., 2018) (top) and *Gabpa* expression in mouse liver from 48h microarray dataset (Hughes et al., 2009) indicated by two different probes.

Motif analysis on 1000bp DNA region surrounding XBP1s peak center did not reveal enriched motifs associated with core circadian clock transcription factors, which is supported by very limited overlap of XBP1s cistrome with those of BMAL1, CLOCK, CRY1, CRY2, NPAS2, PER1, PER2, REV-ERBα, REV-ERBβ and RORα (Figure S6A). Instead, leucine zipper containing transcription factor binding sites, including XBP1, ATF6 and CREB3/CREB3L were strongly enriched as expected (Figure 6B). Both ATF6 and CREB3/CREB3L1/CREB3L2 are known to activate gene expression involved in unfolded protein response (Li et al., 2000; Liang et al., 2006; Tomoishi et al., 2017; Vellanki et al., 2013) and ATF6 and CREB3L2 also exhibit 12h rhythm of gene expression (Figure S6B) (Zhu et al., 2017). These data suggest that ATF6 and CREB3L2 may cooperate with XBP1s in dictating 12h rhythms of gene expression. In addition to leucine zipper transcription factors, we unexpectedly found enriched motifs of ETS transcription factors including GAPB and ELF1 at XBP1s cistrome (Figure 6B). While the ETS transcription factors are known to play important roles in tissue development and cancer progression (Sharrocks, 2001; Sizemore et al., 2017), to the best of our knowledge, their potential involvement in the regulation of CEDIF and crosstalk with XBP1s remain unreported. To determine whether the ETS DNA binding motif mainly localizes mutually exclusive with XBP1 motif or adjacent to the XBP1 motif in *cis*, we re-performed the motif analysis using an 800bp DNA region surrounding the XBP1s peak center, and no longer found the enrichment of these motifs (Figure 6B). This result indicates that GABP and/or ELF1 binding sites mostly occur adjacent to that of XBP1s *in cis* on the same molecule of DNA. GABP transcriptionally regulates gene expression predominantly by forming a heterotetrameric complex composed of two α and two β subunits encoded by the *Gabpa* and *Gabpb1/b2* genes, respectively (Chinenov et al., 2000). By examining a previously published hepatic circadian proteome database (Wang et al., 2018), we found a robust 12h oscillation of nuclear GABPA level (GABPB not reported) (Figure S6C-i). In addition, robust 12h oscillations of nuclear GABPA/GABPB2 (GABPB1 not reported) bound to an artificial DNA fragment harboring GABP consensus sequence were also reported in the same study (Figure S6C-i) (Wang et al., 2018). Both hepatic *Gabpa* and *Gabpb1* mRNA exhibit XBP1s-depenent 12h oscillations (Figures S6C-ii, 7D), and 12h chromatin recruitment of XBP1s to *Gabpb1* gene promoter containing consensus CCACGTCA sequence was also found (Figure S6C-iii). While future efforts are needed to firmly establish the causal roles of GABP in transcriptionally regulating the mammalian 12h rhythms of gene expression, these data, nevertheless, imply the putative concerted actions of GABP with XBP1s in regulating the mammalian 12h-clock (Figure 8A).

**FIGURE 8.**
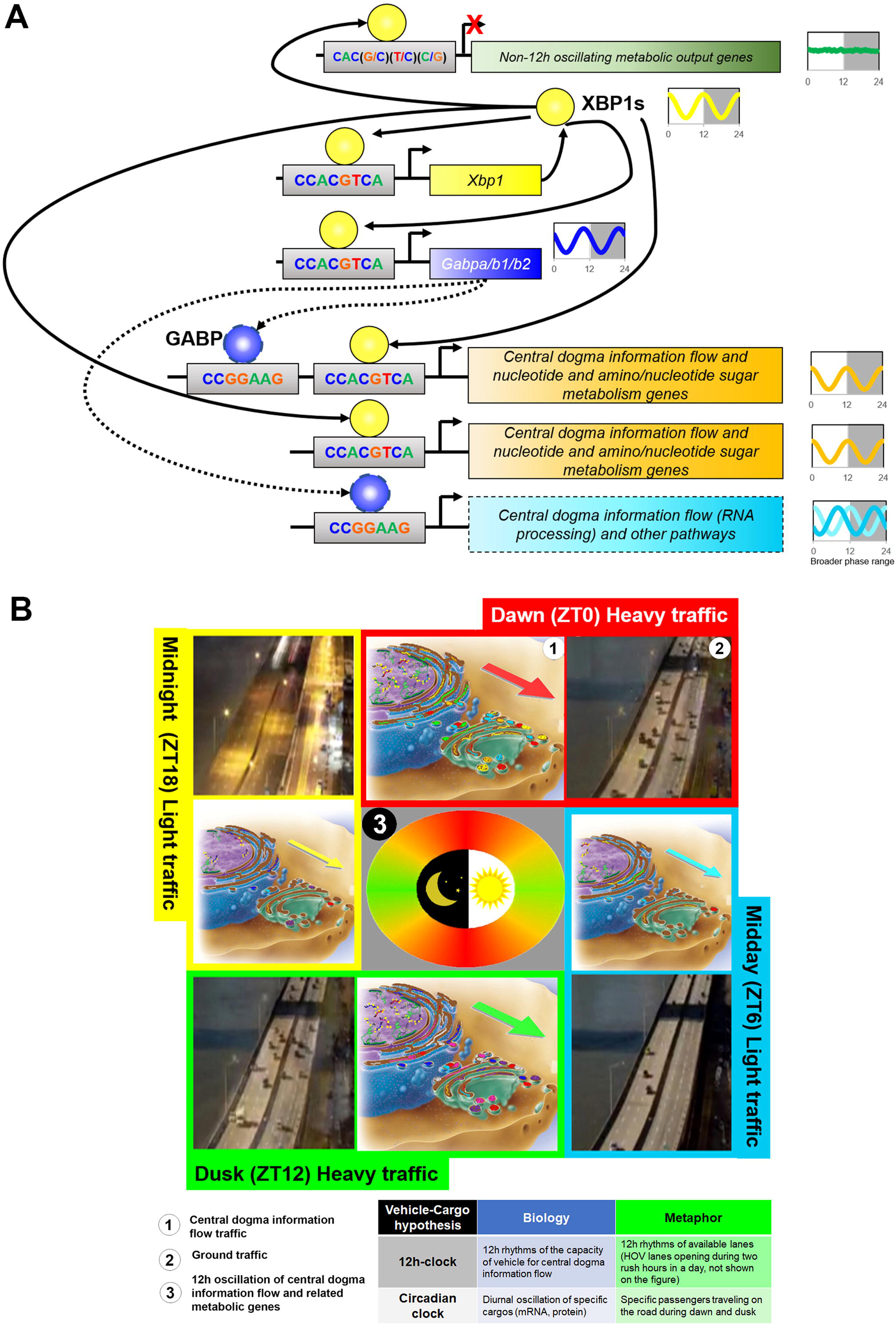
XBP1s transcriptionally regulates 12h rhythms of gene expression involved in central dogma information flow. **(A)** A simplified model summarizing our current understanding of the transcriptional regulation of mammalian 12h-clock by XBP1s. 12h rhythmic XBP1s binding to consensus XBP1s binding motif CCACGCTA within proximal promoter regions drives 12h rhythms of gene expression that are involved in regulating the traffic capacity of CEDIF. XBP1s further self-regulates its own 12h gene expression via this mechanism, thus forming a positive feedforward loop. On the other hand, 12h rhythmic XBP1s binding to degenerate XBP1s binding motif CAC(G/C)(T/C)(C/G) fail to drive 12h rhythms of gene expression that are enriched metabolic pathways. XBP1s further transcriptionally regulates 12h oscillation of GABP transcription factors, whose binding motif exhibits equal strong enrichment on the promoters of XBP1s-dependent 12h genes. Future studies are needed to establish the casual roles of GABP on the transcriptional regulation of 12h-clock (represented by the dashed arrows). **(B)** The vehicle-cargo hypothesis on the distinct functions of 12h-clock versus the 24h circadian clock. Similar to the increased traffic at “rush” hours at each dawn and dusk in people’s daily life, 12h rhythms of CEDIF gene expression peaking at dawn (CT0) and dusk (CT12) (indicated by the circular heat map in the middle) suggest the existence of a 12h oscillation of “traffic” of central dogma information flow (CEDIF), which consists of progressive molecular processing steps from transcription, mRNA processing, ribosome biogenesis, translation, all the way to protein processing/sorting in the ER and Golgi. At this point, it remains to be determined whether it is the total number of molecules undergo processing (illustrated by the varying arrow sizes) or the metabolic rate of processing (not shown in the figure) (or both) that exhibits a 12h oscillation. The vehicle-cargo hypothesis posits: whereas the 12h-clock regulates the 12h-rhythms of the traffic capacity of the CEDIF (thus the vehicle), the circadian clock (and other temporal and tissue-specific mechanisms) contributes to the regulation of diurnal oscillations of specific cargos that undergo molecular processing.

### The motif stringency of XBP1s promoter binding sites dictates XBP1s’ ability to drive 12h-rhythms of transcription of CEDIF genes

Hepatic XBP1s cistrome is predominantly enriched around proximal promoter, compared to enhancer, intragenic and intergenic regions (Figures 6C-E, S6D-F), which is consistent with previously reported XBP1s cistrome distribution in human triple negative breast cancers (Chen et al., 2014). Very intriguingly, we found highly enriched XBP1s binding sites at bidirectional promoter regions (Figures S6E, F). Gene pairs regulated by XBP1s-targeted bidirectional promoters exhibit similar XBP1s-dependent 12h oscillation of expression (Figure S6F). XBP1s transcriptional regulation of bidirectional promoters may reflect a cost-effective way of mammalian 12h-clock control.

We focused our subsequent analysis on proximal promoter XBP1s sites as they can be unambiguously assigned to each gene and thus permit identification of genes exhibiting both XBP1s-dependent 12h-cyling transcriptome and 12h XBP1s cistrome. A limited overlap of these genes was observed, although it reaches statistical significance indicating a dependent relationship between two “omics” (Figure 6F). The 3,431 XBP1s-dependent 12h-cycling genes without XBP1s binding were still enriched in CEDIF and metabolic pathways (Figure S6G). Motif analysis on the promoters of these genes confirmed the lack of XBP1s consensus binding motif but instead revealed strong enrichment of GABP and ELF1 binding sequences (Figure S6H), suggesting these genes are likely transcriptionally regulated by GABP downstream of XBP1s (Figure 8A).

We were surprised to find 679 genes with XBP1s binding near promoters but without 12h transcriptome and eager to identify the potential mechanisms that distinguish these 679 genes (cistrome positive) from the 532 genes (double positive) with both 12h XBP1s cistrome and transcriptome (Figure 6F). GO analysis revealed that while the 532 double positive genes are strongly enriched in CEDIF pathways as expected, the 679 cistrome positive genes completely lack GO terms associated with CEDIF but instead are modestly enriched with various metabolic pathways mainly under circadian clock regulation (Figure 6G). Motif analysis on the promoters of double positive genes reveal expected XBP1s consensus motif CCACGTCA (Figure 6H). Very intriguingly, the top enriched motif for the cistrome positive gene promoters is identified as a *de novo* motif with the sequence CAC(G/C)(T/C)(C/G), which resembles a degenerate XBP1s motif with the 5’ C and 3’ A missing and reduced frequency of the central CACGTC sequences (Figure 6H). XBP1s binding intensity and amplitude to the degenerate motif are weaker compared to consensus XBP1s sequence (Figure 6I). Accordingly, XBP1s binding to the double positive, but not the cistrome positive gene promoters is strongly associated with 12h rhythm of nascent mRNA transcription initiation, as assayed by calculating the area under the curve of pre-pausing portion of Gro-Seq signal (Fang et al., 2014) (Figures 6J-L, S6I, K). Overall, for the 532 double positive genes, we observed a positive correlation among the amplitude of nascent pre-mRNA transcription oscillation (assayed by Gro-Seq), the amplitude of mature mRNA oscillation (assayed by RNA-Seq) and the XBP1s binding motif stringency score (assayed by ChIP-Seq and the higher the score, the more similar the XBP1s binding motif is to the consensus sequence) in XBP1^*Flox*^ mice (Figures 6L-Q, S6J).

Taken together, we hereby demonstrated that the motif stringency of XBP1s binding sites, but not necessarily its expression, dictates its ability to drive 12h rhythms of mRNA transcription: 12h XBP1s chromatin recruitment to gene promoters harboring consensus XBP1s DNA binding motif CCACGTCA drives 12h rhythms of transcription initiation of genes involved in CEDIF and nucleotide/amino sugar metabolism, while weaker binding to promoters with degenerate motifs does not (Figure 8A). Due to the elevated basal level of gene expression among half of the 679 cistrome positive genes in XBP1^*LKO*^ mice (Figures S6K-N), we hypothesize that XBP1s binding to degenerate motif can repress lipid metabolic gene expression possibly through a tethering mechanism by interfering with gene activation by adjacent transcription activators (Herrema et al., 2016).

### The 12h rhythms of CEDIF and nucleotide metabolism gene expression are evolutionarily conserved in marine animals possessing dominant circatidal clock

We originally hypothesized that the mammalian 12h clock evolved from the circatidal clock of coastal and estuarine animals that modulate their behavior in tune to the ∼12.4 hour ebb and flow of the tides (Zhu et al., 2018; Zhu et al., 2017), which were also established independently from the circadian clock (III et al., 2008; Naylor, 1996; Takekata et al., 2012; Takekata et al., 2014; Zhang et al., 2013). To seek further support for our hypothesis, we analyzed two recently published time series RNA-Seq dataset of two marine animals harboring circatidal clock, aposymbiotic sea anemone *Aiptasia diaphaha* (Sorek et al., 2018) and the limpet *Cellana rota* (Schnytzer et al., 2018). In both cases, we found large overlap of 12h-cycling transcritptome between mouse and these two species (Figures 7A-C, S7A-C, Table S12). Computationally constructing predicted interactive network using the overlapping 12h-cycling genes in both species via STRING (Szklarczyk et al., 2015) or by traditional GO analysis revealed sub-hubs involved in different steps of CEDIF and nucleotide metabolism (Figures 7B and S7B, Table S13). We further observed 12h oscillations of *Gabpa* in both *Aiptasia diaphaha* and *Cellana rota* and *Gabpb2* in *Aiptasia diaphaha* (Figures 7D and S7D). 12h oscillation of *Xbp1* in *Cellana rota* was previously reported (Zhu et al., 2018). These data combined with the recent independent report of earlier evolutionary origin of 12h-cycling genes than circadian genes (Castellana et al., 2018) provides strong support for our hypothesis that the mammalian 12h clock evolved from the circatidal clock.

## DISCUSSION

In this section, we will mainly discuss two aspects of the mammalian 12h-clock, its transcriptional regulation and physiological functions.

### The working model of the transcriptional regulation of mammalian 12h-clock by XBP1s

Our study establishes XBP1s as a master transcriptional regulator of the mammalian 12h-clock. Hepatic ablation of XBP1 results in the impairment of 87% of hepatic 12h-cycling transcriptome. XBP1s directly transcriptionally regulates more than 500 genes (with acrophases around CT2 and CT14) via rhythmic binding to promoters containing consensus sequence CCACGTCA. Alternatively, XBP1s transcriptionally regulates 12h oscillations of GABP expression, which in turn can bind to gene promoters harboring ETS consensus sequence CCGGAAG and putatively regulates additional 12h transcriptome with a wider range of acrophases. GABP can also act in concert with XBP1s *in cis* on a subset of genes containing adjoining XBP1s and GABP DNA binding motif in gene promoters (Figure 8A). XBP1s also regulates its own 12h transcription, therefore completing a positive feedforward loop (Figure 8A) (Majumder et al., 2012). At this point, what remains elusive is the mechanism(s) of negative feedback required for sustaining cell-autonomous oscillations of the 12h-clock. One potential candidate is the unspliced form of XBP1 (Xbp1us), which has been previously shown to negatively regulate UPR by forming a complex with XBP1s and ATF6 in the cytosol and target them for ubiquitin-mediated degradation (Yoshida et al., 2006; Yoshida et al., 2009). Supporting this hypothesis is the observed 12h rhythm of *Xbp1us* mRNA (data not shown), although more evidence is needed in the future. One last note, what is summarized in Figure 8 is the current working model of the transcriptional regulation of mammalian 12h-clock by XBP1s, it is subject to revision and modification with more available experimental and mathematical modeling data in the future.

### The vehicle-cargo hypothesis on the distinct functions of 12h-clock versus the circadian clock

A very fascinating finding from our study is the coordinated 12h oscillations of genes involved in the entire central dogma information flow, ranging from mRNA transcription, mRNA processing, translation regulation to protein processing and sorting in the ER and Golgi, which include both anabolic and catabolic processes. The vast majority of these genes peak at dawn (CT0-CT3) and dusk (CT12 to CT15), corresponding to the transition periods between fasting/feeding and rest/activity that are associated with elevated metabolic stress (Zhu et al., 2018). In light of these findings, we hereby propose a vehicle-cargo hypothesis that attempts to decipher the distinct functions of 12h-clock versus the circadian clock (Figure 8B). We argue that the 12h-clock accommodates rush hours’ (at dawn and dusk) elevated gene expression and processing by controlling the 12h rhythms of the global traffic capacity (and/or the traffic rate) of the central dogma information flow (thus the vehicle), in tune to the 12h cycle of metabolic stress (Zhu et al., 2018) (Figure 8B). The circadian clock and/or other tissue specific pathways, on the other hand, dictate the particular genes/gene products processed at each rush hour (thus the cargo) (Figure 8B). An everyday metaphor would be the fluctuating daily traffic on the highway: the 12h-clock is comparable to the highway that increases its operating capacity during the morning and evening rush hours (by opening the HOV lane, for example), while the 24h circadian clock dictates the specific cars that go on the highway during morning and evening (Figure 8B). Future efforts should be directed toward characterizing the temporal profile of 12h-clock-dependent mRNA and protein being processed in the nucleus and ER/Golgi, respectively.

## MATERIALS AND METHODS

### Cell Lines

Mouse embryonic fibroblast (MEF) was prepared as previously described (Stashi et al., 2014).

### Animals

XBP1^Flox^ mice was previously described (Lee et al., 2008). XBP1^*LKO*^ mice were generated by crossing Albumin-CRE mice with XBP1^Flox^ mice. All mice are in C57BL/6 background, male and between 3 and 4 months of age. Mice were first entrained under LD12:12 conditions for 2 weeks before transferred to constant darkness for 24hrs. Mice were then sacrificed at a 2h interval for a total of 48 hrs. Mice were fed *ad libitum* during the entire experiment. The Baylor College of Medicine and University of Pittsburgh Institutional Animal Care and Utilization Committee approved all experiments.

### Food Intake Monitoring

Comprehensive Lab Animal Monitoring System (CLAMS) Calorimetry (Columbus Instruments) was used for real-time measuring of food intake. XBP1^Flox^ (n=4) and XBP1^*LKO*^ (n=4) mice were acclimated to the chambers for at least one week and ad libitum food intake was monitored for 72 hours under LD12:12 followed by 96 hours of constant darkness.

### Locomotor Activity Monitoring

The Home Cage Activity System (Omnitech Electronics, Inc) was used for real-time measuring of spontaneous locomotor activity in a home-cage environment. XBP1^Flox^ (n=4) and XBP1^*LKO*^ (n=4) mice were acclimated to the home cage for at least one week and fed ad libitum. Spontaneous locomotor activity was measured under either LD12h:12h condition or constant darkness conditions.

### siRNA Transient Transfections

MEFs were transfected with 10µM of different siRNAs for 24∼48 hours with Lipofectamine RNAiMAX reagents (Life technologies) per the manufacturer’s instructions. Source of siRNA are as follows: siGENOME Non-Targeting siRNA pool (Dharmacon, D-001206-1305), siGENOME SMARTpool ARNTL (Dharmacon, L-040483-01-0005), siGENOME SMARTpool XBP1 (Dharmacon, L-040825-00-0005).

### Synchronization of MEFs

MEFs were isolated from male SRC-2^*fl/fl*^ mice and immortalized by SV40 T antigen as previously described (Stashi et al., 2014). For tunicamycin treatment, MEFs were cultured in DMEM (4.5g/L glucose) supplemented with 10% FBS and treated with 25ng/ml of tunicamycin for 2h, and then washed with 1X PBS before cultured in the same medium. For dexamethasone treatment, MEFs were cultured in DMEM (4.5g/L glucose) supplemented with 10% FBS and treated with 100nM Dex for 30mins, and then washed with 1X PBS before cultured in the same medium. For all cell culture experiments, cells were cultured at 37 °C with 5% CO2.

### Immunoblot

Immunoblot analyses were performed as described previously (Zhu et al., 2015). Briefly, proteins separated by 4∼20% gradient SDS-PAGE gels (Biorad) were transferred to nitrocellulose membranes, blocked in TBST buffer supplemented with 5% bovine serum albumin (BSA) and incubated overnight with primary anti-XBP1s antibody (Biolegend Poly6195) at 4°C. Blots were incubated with an appropriate secondary antibody coupled to horseradish peroxidase at room temperature for 1 hour, and reacted with ECL reagents per the manufacturer’s (Thermo) suggestion and detected on X-ray film by autoradiography.

### qRT-PCR

Total mRNA was isolated from murine embryonic fibroblasts (MEFs) with PureLink RNA mini kit (Life Technologies) per the manufacturer’s instructions. Reverse transcription was carried out using 5µg of RNA using Superscript III (Life Technologies) per the manufacturer’s instructions. For gene expression analyses, cDNA samples were diluted 1/30-fold (for all other genes except for 18sRNA) and 1/900-fold (for 18sRNA). qPCR was performed using the Taqman or SYBR green system with sequence-specific primers and/or the Universal Probe Library (Roche). All data were analyzed with 18S or β*-actin* as the endogenous control. qPCR primer sequences are as follows:

Mouse total *Xbp1* forward primer: gggtctgctgagtcc

Mouse total Xbp1 reverse primer: cagactcagaatctgaagagg

Mouse *Xbp1s* forward primer: ccgcagcaggtgc

Mouse *Xbp1s* reverse primer: cagactcagaatctgaagagg

Mouse *Xbp1us* forward primer: actatgtgcacctctgcag

Mouse *Xbp1us* reverse primer: cagactcagaatctgaagagg

Mouse *Arntl* forward primer: gccccaccgacctactct

Mouse *Arntl* reverse primer: tgtctgtgtccatactttcttgg

Mouse *Nr1d1* forward primer: acgaccctggactccaataa

Mouse *Nr1d1* reverse primer: ccattggagctgtcactgtaga

Mouse *Per2* forward primer: caacacagacgacagcatca

Mouse *Per2* reverse primer: tcctggtcctccttcaacac

Mouse *Ero1lb* forward primer: atgattcgcaggaccacttt

Mouse *Ero1lb* reverse primer: tcagcagcaggtccacatac

Mouse *Paics* forward primer: tagcactccagaggctgagg

Mouse *Paics* reverse primer: ctccacggcaagttgagtc

Mouse *Srp72* forward primer: ccaaagcgtgtctaatcctga

Mouse *Srp72* reverse primer: ggtcaccaatgcagatacca

Mouse *Sf3a1* forward primer: gatgatgaggtttatgcaccag

Mouse *Sf3a1* reverse primer: agtacgtcgctcagccaact

Mouse *Tmed7* forward primer: agttggagaagacccaccttt

Mouse *Tmed7* reverse primer: agagcttcatgaatggaaacg

Mouse 18s RNA forward primer: ctcaacacgggaaacctcac

Mouse 18s RNA reverse primer: cgctccaccaactaagaacg

Mouse *β-actin* forward primer: aaggccaaccgtgaaaagat

Mouse *β-actin* reverse primer: gtggtacgaccagaggcatac

### RNA-Seq

Mouse liver tissues were collected from XBP1^Flox^ (n=2) and XBP1^*LKO*^ (n=2) mice at 2h interval for a total of 48 hours under constant darkness condition. Total RNA was isolated from mouse liver with miRNeasy Mini Kit (Qiagen) per the manufacturer’s instructions. Extracted RNA samples underwent quality control assessment using the RNA Pico 6000 chip on Bioanalyzer 2100 (Agilent) and were quantified with Qubit Fluorometer (Thermo Fisher). Strand-specific total mRNA-Seq libraries were prepared using the Universal Plus mRNA-Seq kit (NuGen) per the manufacturer’s instructions using 200ng of RNA. The size selection for libraries were performed using SPRIselect beads (Beckman Coulter) and purity of the libraries were analyzed using the High Sensitivity DNA chip on Bioanalyzer 2100 (Agilent). The prepared libraries were pooled and sequenced using NoveSeq 6000 (Illumina), generating ∼20 million 2×100 bp paired-end reads per samples. RNA-Seq library preparation and sequencing were performed at University of Houston Seq-N-Edit Core. Reads were mapped to the mouse genome build mm10 using HISAT (Kim et al., 2015), and gene expression was normalized and quantified using StringTie (Pertea et al., 2015) for FPKM values using default parameters in Python.

### Identification of Oscillating Transcriptome

Averaged FPKM values at each time were used for cycling transcripts identification. To identify cycling transcriptome, we first filtered out transcripts that exhibit an average FPKM value below 0.01 and 25,289 transcripts remain. Superimposed oscillations for all 25,289 transcripts were identified using previously described eigenvalue/pencil method (Antoulas et al., 2018; Zhu et al., 2017). Specifically, three oscillations were identified from each gene. Criterion for circadian genes are: period between 22.5h to 25.5h, decay rate between 0.8 and 1.2 and mean expression FPKM larger than 0.1; for ∼12h genes: period between 10.5h to 13.5h, decay rate between 0.8 and 1.2 and mean expression FPKM larger than 0.1; for ∼8h genes: period between 7h to 9h, decay rate between 0.8 and 1.2 and mean expression FPKM larger than 0.1. To determine the FDR of identification of rhythmic transcripts, we used a permutation-based method that randomly shuffles the time label of gene expression data and subject each of the permutation dataset to eigenvalue/pencil method applied with the same criterion (Rey et al., 2018). These permutation tests were run 1,000 times, and FDR was estimated by taking the ratio between the mean number of rhythmic profiles identified in the permutated samples (false positive ones) and the number of rhythmic profiles identified in the original data. All the analysis were performed in MatlabR2017A. Heat maps were generated by Gene Cluster 3.0 and TreeView 3.0 alpha 3.0 using log2 mean-normalized values.

### T-distributed Stochastic Neighbor Embedding (***t*-SNE) ANALYSIS**

t-SNE analysis was performed on identified pure 12h oscillations using MatlabR2017A. “Exact’ algorithm and “city block” distance metric were used.

### Chromatin Immunoprecipitation (ChIP)-Seq

ChIP for XBP1s was performed using anti-XBP1s antibody (Biolegend Poly6195) as previously described (Zhu et al., 2015). Briefly, mouse liver samples were submerged in PBS + 1% formaldehyde, cut into small (∼1 mm3) pieces with a razor blade and incubated at room temperature for 15 minutes. Fixation was stopped by the addition of 0.125 M glycine (final concentration). The tissue pieces were then treated with a TissueTearer and finally spun down and washed twice in PBS. Chromatin was isolated by the addition of lysis buffer, followed by disruption with a Dounce homogenizer. The chromatin was enzymatically digested with MNase. Genomic DNA (Input) was prepared by treating aliquots of chromatin with RNase, Proteinase K and heated for reverse-crosslinking, followed by ethanol precipitation. Pellets were resuspended and the resulting DNA was quantified on a NanoDrop spectrophotometer. An aliquot of chromatin (30 μg) was precleared with protein A agarose beads (Invitrogen). Genomic DNA regions of interest were isolated using 4 μg of antibody. Complexes were washed, eluted from the beads with SDS buffer, and subjected to RNase and proteinase K treatment. Crosslinking were reversed by incubation overnight at 65 °C, and ChIP DNA was purified by phenol-chloroform extraction and ethanol precipitation. The DNA libraries were prepared and sequenced at University of Houston Seq-N-Edit Core per standard protocols. DNA libraries were prepared with Ovation^®^ Ultralow V2 DNA-Seq library preparation kit (NuGen) using 1ng input DNA. The size selection for libraries were performed using SPRIselect beads (Beckman Coulter) and purity of the libraries were analyzed using the High Sensitivity DNA chip on Bioanalyzer 2100 (Agilent). The prepared libraries pooled and sequenced using NextSeq 500 (Illumina), generating ∼15 million 76 bp single-end reads per samples.

### ChIP-Seq analysis

Duplicates were pooled at each time for subsequent ChIP-Seq analysis. The sequences identified were mapped to the mouse genome (UCSC mm10) using BOWTIE function in Galaxy. Only the sequences uniquely mapped with no more than 2 mismatches were kept and used as valid reads. PCR duplicates were also removed. Peak calling was carried out by MACS (version 1.4.2 20120305) in Galaxy/Cistrome (options --mfold 10, 30 --pvalue 1e-4), on each ChIP-seq file against the ChIP-Seq of XBP1^*LKO*^ mice. To account for the different sequencing depths between samples, the signal files generated from MACS were RPKM normalized to sequencing depth (Meyer and Liu, 2014). Due to the robust 12h oscillation of XBP1s hepatic nuclear proteins (Cretenet et al., 2010; Zhu et al., 2017), only XBP1s cistrome exhibiting robust 12h oscillations (period between 10.5h to 13.5h; decay rate between 0.8 to 1.2; phase between 0h to 3h) are selected as bona fide XBP1s binding sites.

### Gene Ontology Analysis

DAVID (Huang da et al., 2009) (https://david.ncifcrf.gov) was used to perform Gene Ontology analysis. Briefly, gene names were first converted to DAVID-recognizable IDs using Gene Accession Conversion Tool. The updated gene list was then subject to GO analysis using Mus musculus as background and with Functional Annotation Chart function. KEGG_PATHWAY, were used as GO categories for all GO analysis. Only GO terms with p value smaller than 0.05 were included for further analysis.

### Binding Site Annotation and Profiling

CEAS (Cis-regulatory Element Annotation System) function in Galaxy/Cistrome was applied to calculate the enrichment of the binding sites in the promoter, exon, intron, UTR and other genomic regions against the mappable mouse genome using the binomial model.

### Motif Analysis

Motif analysis was performed with Discriminative DNA Motif Discovery (DREAM) tool (version 4.9.1) or the SeqPos motif tool (version 0.590) embedded in Galaxy Cistrome.

### Network Analysis

Construction of interacting network of evolutionarily conserved 12h genes were performed by STRING (https://string-db.org/).

### *Post-hoc* **Analysis of Serum Synchronized MMH-D3 Transcriptome**

The time series transcriptome of serum-synchronized murine liver MMH-D3 cell was published previously (Atwood et al., 2011). Transcripts with averaged expression larger than 20 were used for subsequent analysis. Upon examining the raw data, we observed noticeable baseline changes in most reported mRNA oscillations. Therefore, we subjected the raw data to polynomial detrend (n=3) [we found higher order of polynomial detrend (n>3) will lead to the disappearance of circadian rhythm in some core circadian genes, therefore we think n=3 is the optimal trade-off between over-fitting and under-fitting]. The polynomial detrended data were then subject to eigenvalue/pencil analysis to identify superimposed oscillations. Specifically, three oscillations were identified from each gene. Criterion for circadian genes are: period between 20.5h to 23.5h, decay rate between 0.9 and 1.1; for ∼12h genes: period between 9.5h to 12.5h, decay rate between 0.9 and 1.1. The smaller periods for both circadian and 12h genes were selected based upon the distribution of all period uncovered as shown in **Figure S5A**.

## DATA ACCESS

All raw and processed sequencing data generated in this study have been submitted to the NCBI Gene Expression Omnibus (GEO; http://www.ncbi.nlm.nih.gov/geo/) under accession number GSE126335.

## ACKNOWLEDGEMENTS

We want to thank Drs. Brian York, and Maricarmen Delia Planas-Silva for assistance with equipment and experiments. This research was supported by CPRIT grant RP170005 to C.C. and the American Diabetes Association junior faculty development award 1-18-JDF-025 to B.Z.

## SUPPLEMENTAL FIGURE

**FIGURE S1.**
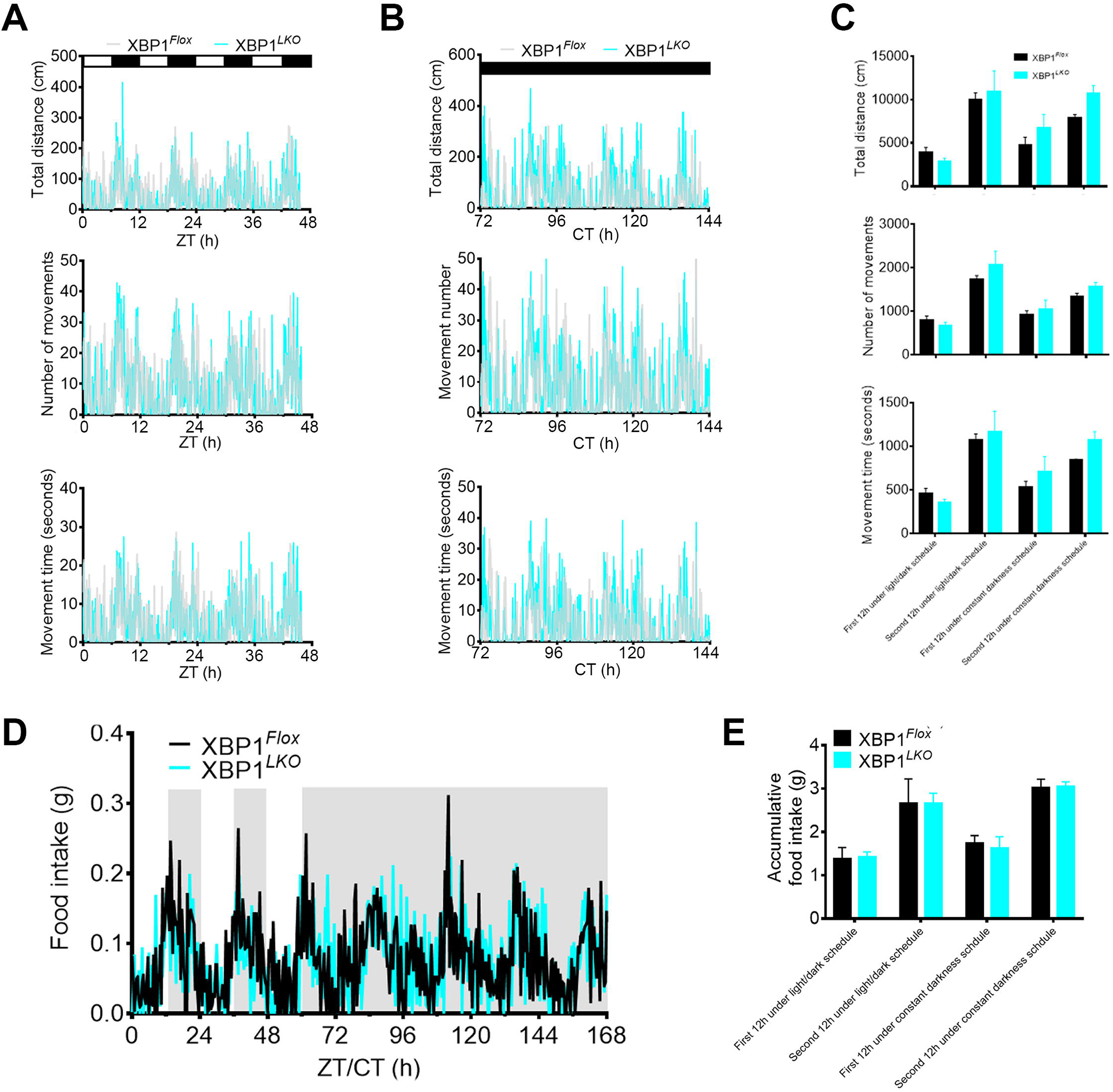
Liver-specific deletion of XBP1s does not alter rhythmic locomotor activity nor fasting-feeding cycles in mice, related to Figure 1. **(A, B)** Real-time home cage activity monitoring of total distance covered (top), number of movements recorded (middle) and movement time recorded (bottom) in XBP1^*Flox*^ and XBP1^*LKO*^ mice under 12h light/12h dark conditions **(A)** and constant darkness condition. **(B). (C)** Averaged measurements within the first and second 12h of a day as described in **A** and **B**. **(D)** Real-time measurement of food intake in XBP1^*Flox*^ and XBP1^*LKO*^ mice under both 12h light/12h dark and constant darkness condition measured by CLAMS system. **(E)** Averaged measurements within the first and second 12h of a day as described in **D**. Data are graphed as the mean ± SEM (n = 4).

**FIGURE S2.**
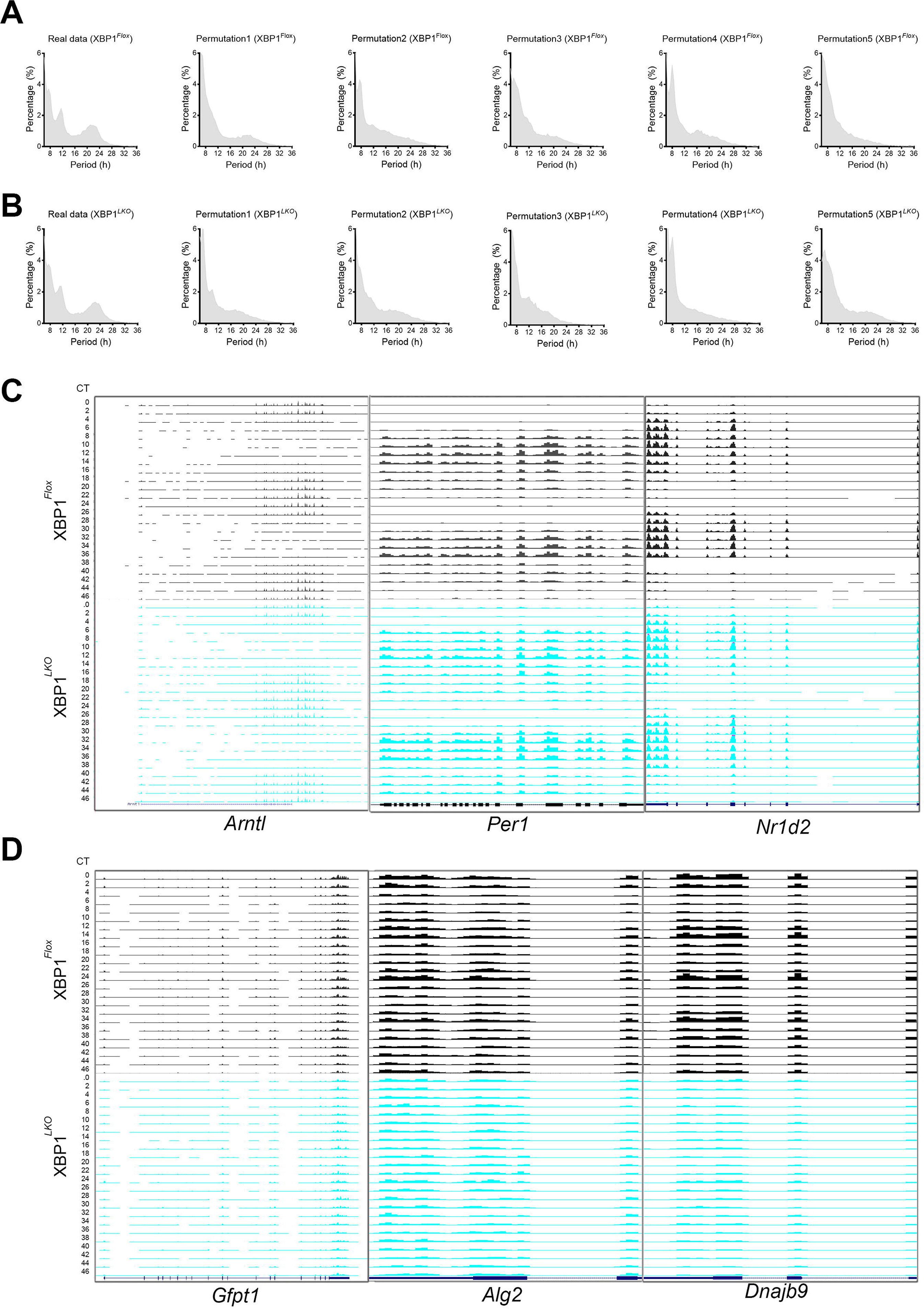
Liver-specific deletion of XBP1s impairs global hepatic 12h-transcriptome, but not the circadian rhythm in mice, related to Figure 1. **(A, B)** Permutation was performed on the raw data by randomly shuffling the time label. Distribution of periods of all oscillations identified by the eigenvalue/pencil method from five representative permutated dataset from XBP1^*Flox*^ mice (**A**) and XBP1^*LKO*^ mice (**B**). **(C, D)** UCSC genome browser snapshot view of RNA-Seq tracks of selective circadian **(C)** and 12h-cycling **(D)** gene expression in XBP1^*Flox*^ mice and XBP1^*LKO*^ mice.

**FIGURE S3.**
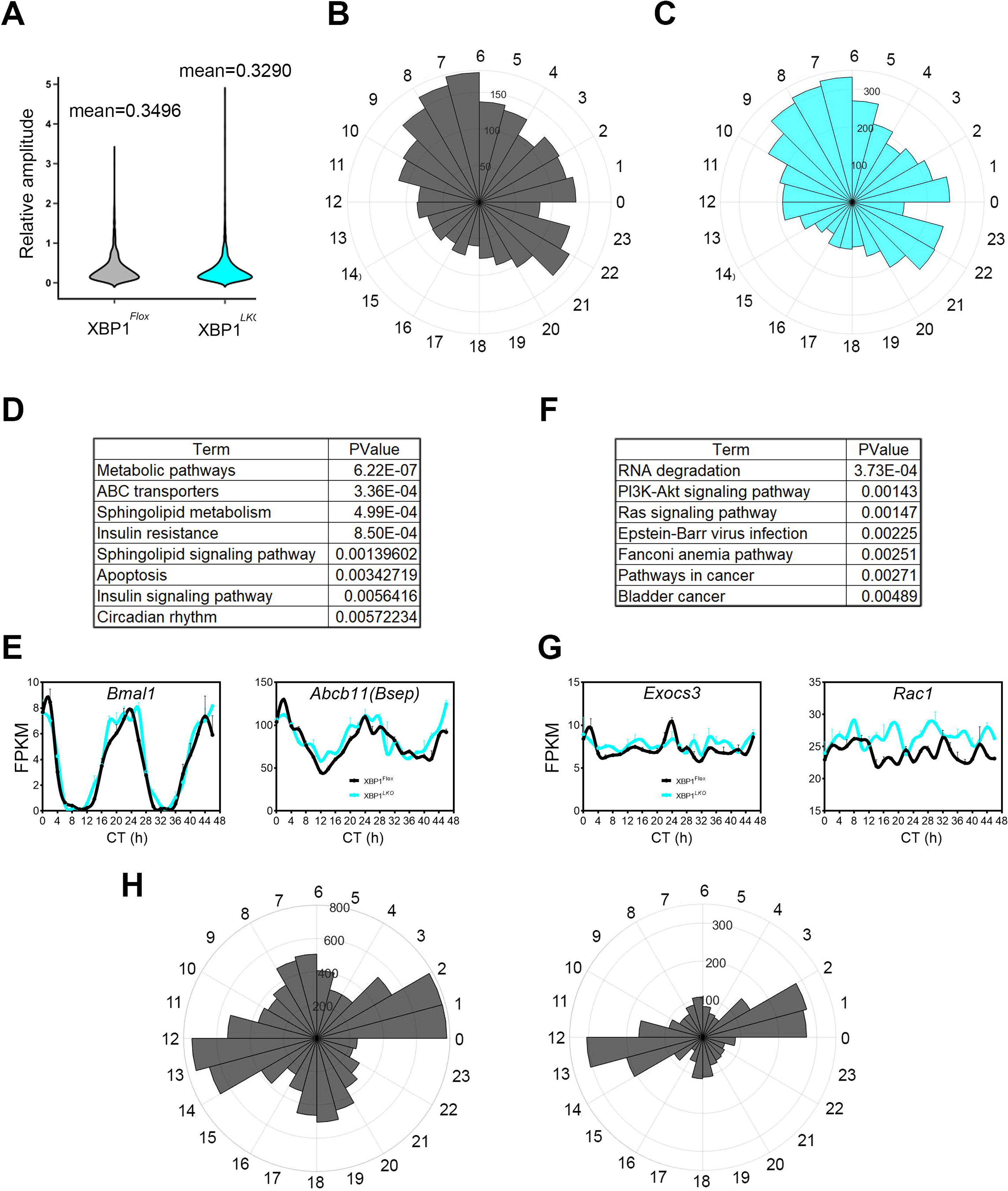
The effects of liver-specific deletion of XBP1s on hepatic circadian gene expression, related to Figure 1. **(A)** Relative amplitudes of all superimposed 24h rhythms from XBP1^*Flox*^ mice and XBP1^*LKO*^ mice identified by the eigenvalue/pencil method. **(B, C)** Polar histograms demonstrating phase distributions of all superimposed 24h rhythms from XBP1^*Flox*^ **(B)** and XBP1^*LKO*^ **(C)** mice. **(D, E)** 1,763 genes with superimposed 24h rhythms found in both XBP1^*Flox*^ and XBP1^*LKO*^ mice. GO analysis showing enriched KEGG pathways and their corresponding p values **(D)** and RNA-Seq data for representative genes **(E) (F, G)** 798 genes with superimposed 24h rhythms only found in XBP1^*Flox*^ mice. GO analysis showing enriched KEGG pathways and their corresponding p values **(F)** and RNA-Seq data for representative genes **(G). (H)** Polar histograms demonstrating phase distributions of all (left) and dominant (right) 12h rhythms identified from XBP1^*Flox*^ mice.

**FIGURE S4.**
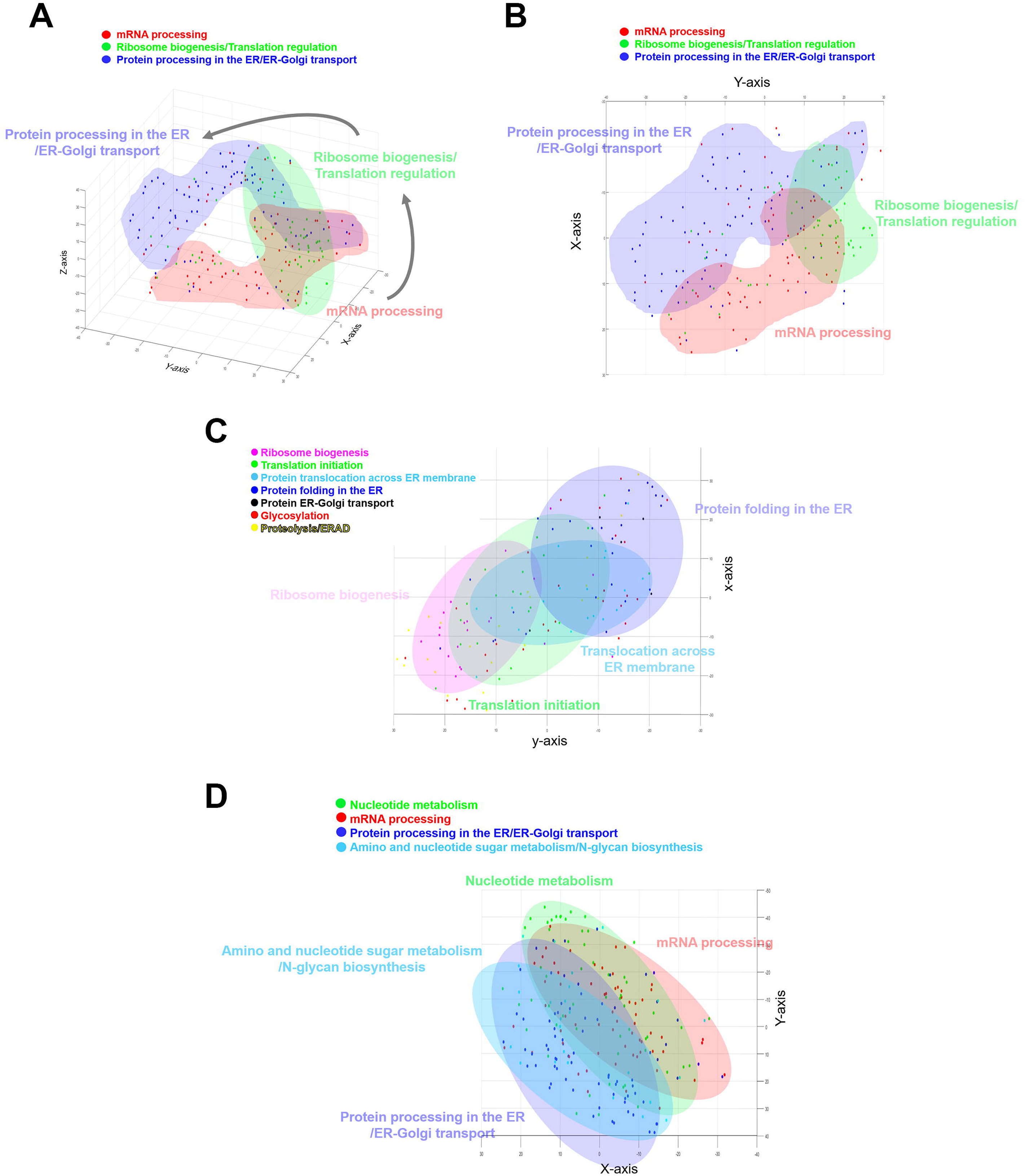
Clustering CEDIF genes based upon their 12h transcriptome profile reveals a spatial order consistent with the direction of central dogma information flow, related to Figures 3, 4. **(A, B)** Clustering of genes involved in mRNA processing, ribosome biogenesis/translation initiation and protein processing and transport based upon their superimposed 12h transcriptome projected onto 3D (viewed from a different angle compared to Figure 3B) **(A)** and 2D **(B)** t-SNE space. **(C)** Clustering of genes involved in different sub-steps of protein metabolism based upon their superimposed 12h transcriptome projected onto 2D t-SNE space. **(D)** Clustering of genes involved in different metabolic pathways and CEDIF based upon their superimposed 12h transcriptome projected onto 2D t-SNE space.

**FIGURE S5.**
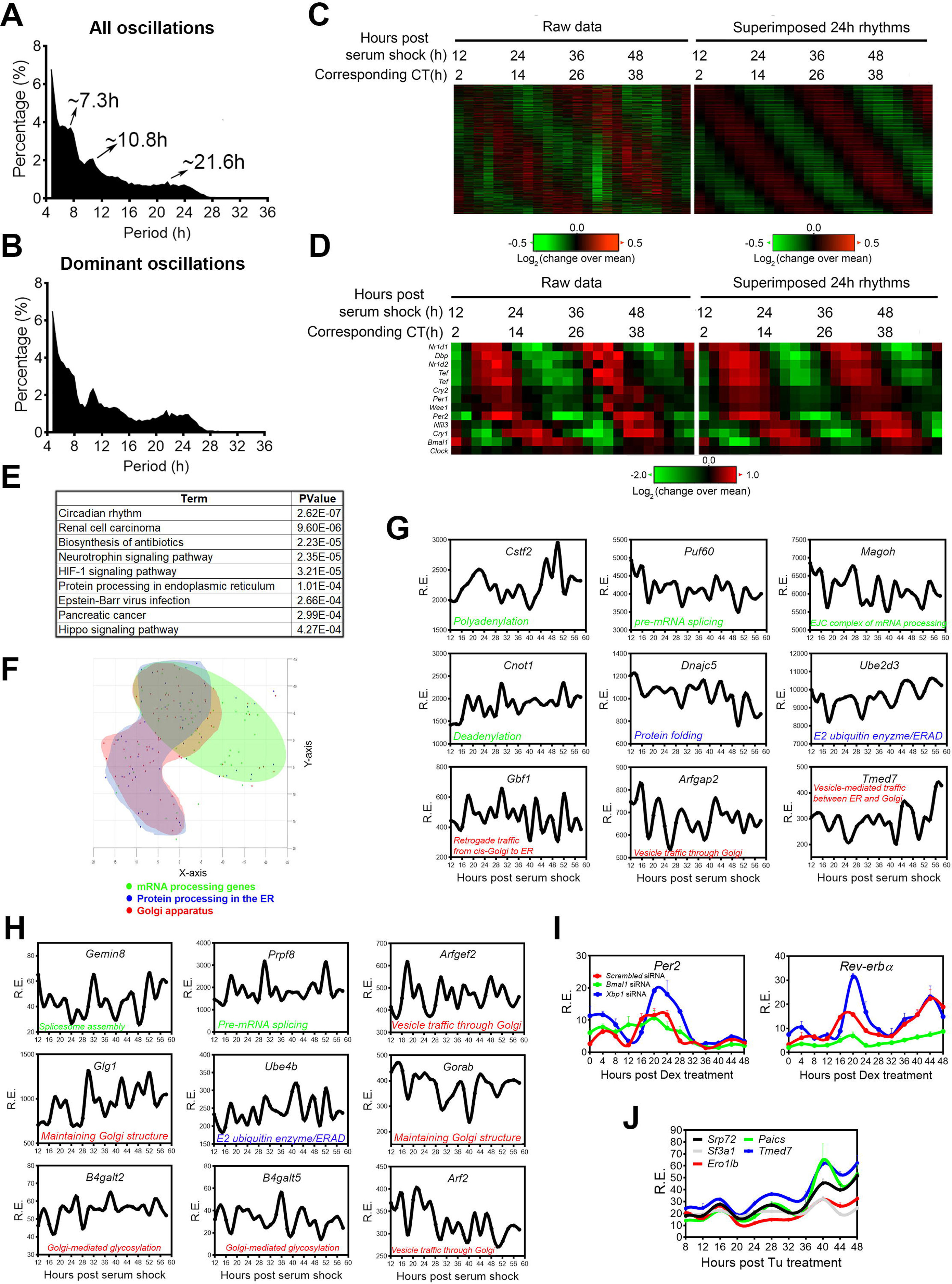
The 12h-rhythms of CEDIF gene expression are cell-autonomous, related to Figure 5. **(A, B)** Distribution of periods of all **(A)** and dominant oscillations **(B)** identified by eigenvalue/pencil method from MMH-D3 cells. **(C, D)** Heat map of all circadian **(C)** and core circadian clock **(D)** gene expression identified from MMH-D3 cells with both raw data and superimposed 24h rhythms shown. Both the original time after serum shock, as well as converted time in CT are shown. **(E)** GO analysis showing enriched KEGG pathways and their corresponding p values for all circadian gene identified in **C**. **(F)** Clustering of MMH-D3 genes involved in different CEDIF based upon their superimposed 12h transcriptome projected onto 2D t-SNE space **(G)** Microarray data of representative ∼12h-cycling genes involved in CEDIF in MMH-D3 cells that are commonly shared with mouse liver. **(H)** Microarray data of representative ∼12h-cycling genes involved in CEDIF uniquely found in MMH-D3 cells, but not in mouse liver. **(I)** MEFs were transfected with different siRNAs and treated with Dexamethasone (100nM) for 30min and qPCR was performed at different time points post Dex shock. **(J)** MEFs were treated with Tu (25ng/µl) for 2hr and qPCR was performed at different time points post Tu shock.

**FIGURE S6.**
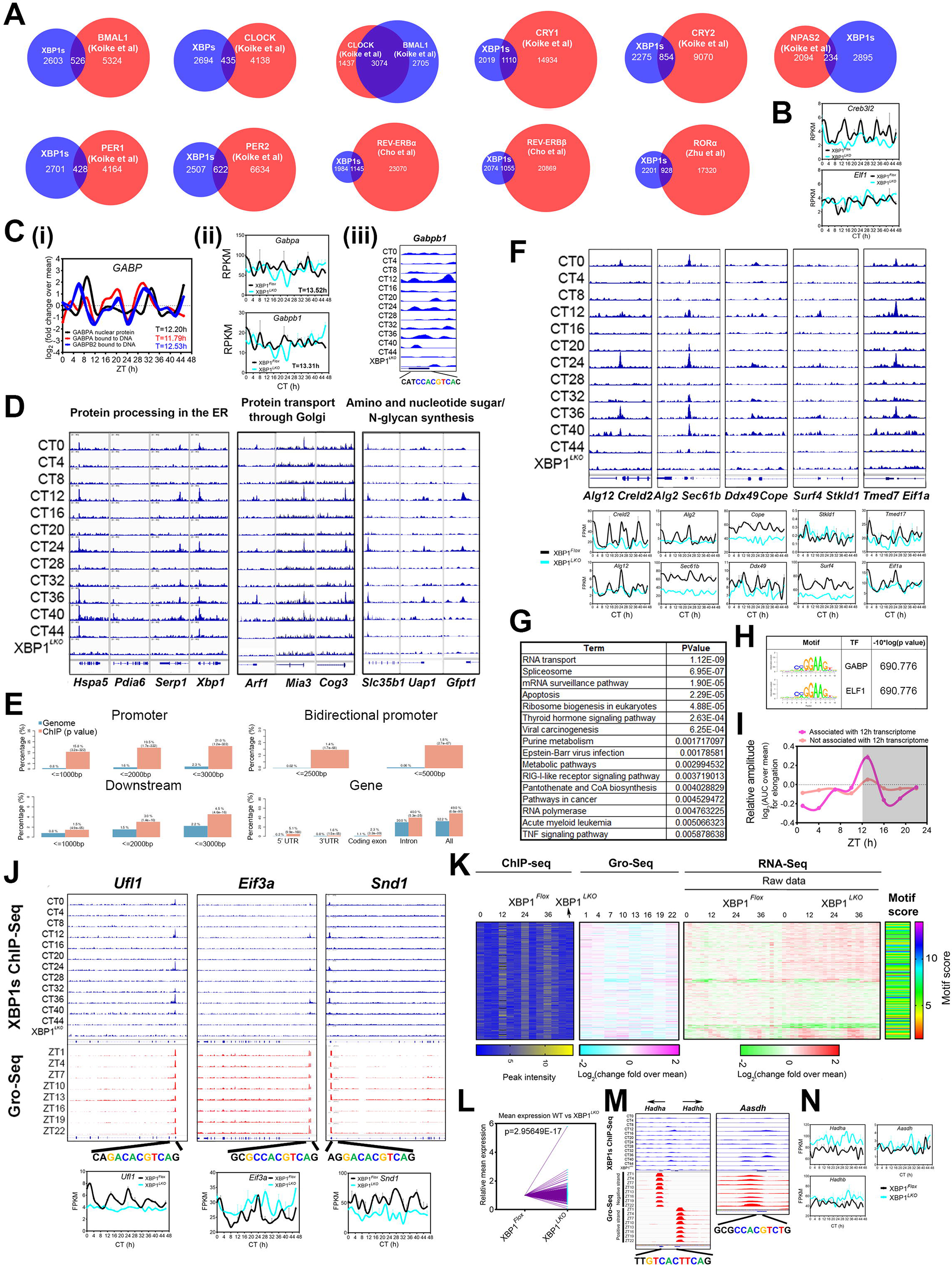
The motif stringency of XBP1s promoter binding sites dictates XBP1s’ ability to drive 12h-rhythms of transcription of CEDIF genes, related to Figure 6. **(A)** Venn diagram depicting common and unique XBP1s cistrome and those of core circadian clock transcription factors compiled from (Cho et al., 2012; Koike et al., 2012; Zhu et al., 2015). **(B)** RNA-Seq data for *Creb3l2* and *Elf1* in XBP1^*Flox*^ and XBP1^*LKO*^ mice. **(C)** 12h rhythms of GABP expression regulated by XBP1s. **(i)** Total nuclear level of GABPA and nuclear level of GABPA and GABPB2 bound to DNA compiled from (Wang et al., 2018). **(ii)** RNA-Seq data for *Gabpa* and *Gabpb1* in XBP1^*Flox*^ and XBP1^*LKO*^ mice with calculated periods in XBP1^*Flox*^ mice shown. **(iii)** Snapshot of *Gabpb1* promoter for alignment of XBP1s binding sites at different CTs in liver tissue of XBP1^*Flox*^ and XBP1^*LKO*^ mice, with XBP1s consensus motif also shown. **(D)** Snapshot of selected genes for alignment of XBP1s binding sites at different CTs in liver tissue of XBP1^*Flox*^ and XBP1^*LKO*^ mice. **(E)** Percentage of XBP1s cistromes located at different positions relative to target genes compared with that of mouse genome. **(F)** Snapshot of selected genes for alignment of XBP1s binding sites at bidirectional promoters at different CTs in liver tissue of XBP1^*Flox*^ and XBP1^*LKO*^ mice and RNA-Seq data for these genes in XBP1^*Flox*^ and XBP1^*LKO*^ mice. **(G)** GO analysis showing enriched KEGG pathways and their corresponding p values for 3,431 12h transcriptome without XBP1s binding sites. **(H)** Top enriched SeqPos motifs common to proximal promoters (600bp around TSS) of 3,431 12h genes without XBP1s binding sites. **(I)** Log_2_ mean normalized transcription elongation rates calculated from the Gro-Seq data (Fang et al., 2014) for XBP1s target genes with or without associated 12h transcriptome. **(J)** Snapshot of target genes selected for alignment of XBP1s binding sites at different CTs in liver tissue of XBP1^*Flox*^ and XBP1^*LKO*^ mice as well as published Gro-Seq data (Fang et al., 2014). Consensus XBP1s binding motifs located at each gene promoter are also shown. **(K-N)** 679 genes with proximal promoter XBP1s binding but no 12h transcriptome in XBP1^*Flox*^ mice. **(K)** Comparisons of heat maps of XBP1s binding intensity, transcription initiation rates calculated from Gro-Seq (Fang et al., 2014), gene expression in XBP1^*Flox*^ and XBP1^*LKO*^ mice, and XBP1s binding motif score. **(L)** Average expression for each gene in XBP1^*Flox*^ and XBP1^*LKO*^ mice. **(M)** Snapshot of target genes selected for alignment of XBP1s binding sites at different CTs in liver tissue of XBP1^*Flox*^ and XBP1^*LKO*^ mice as well as published Gro-Seq data (Fang et al., 2014). Degenerate XBP1s binding motifs located at each gene promoter are also shown. **(N)** RNA-Seq data for representative genes in XBP1^*Flox*^ and XBP1^*LKO*^ mice. Data are graphed as mean ± SEM (n=2).

**FIGURE S7.**
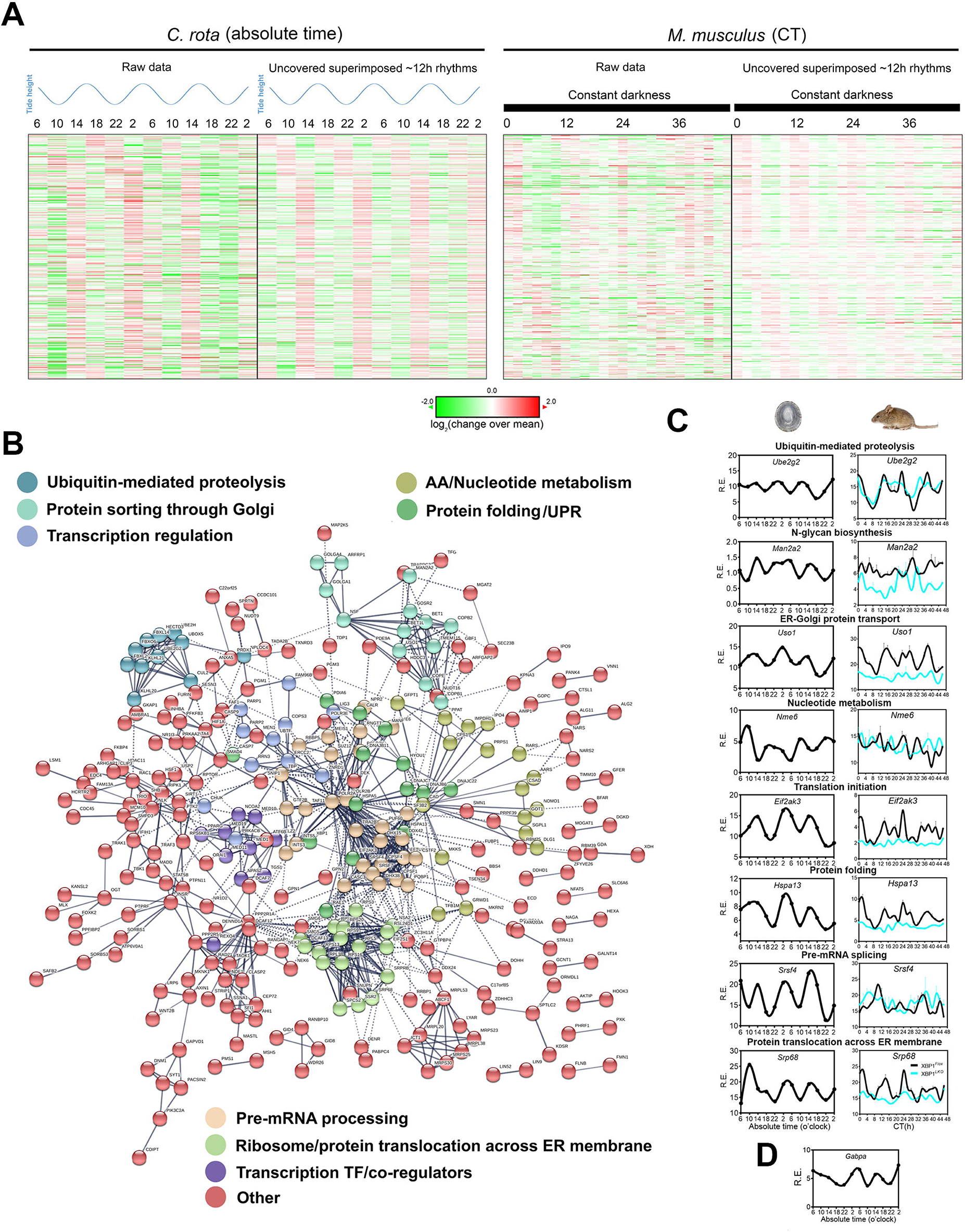
The 12h-rhythms of CEDIF gene expression are evolutionarily conserved in the limpet *C. rota*, which possesses a dominant circatidal clock, related to Figure 7. **(A)** Heat map of side-by-side comparison of evolutionarily conserved 12h gene expression in *C. rota* (Schnytzer et al., 2018) and mouse liver, with both raw data and superimposed ∼12h rhythms shown. The level of tides corresponding to each time was also shown. **(B)** Predicted interactive network construction of these conserved 12h-cycling genes using STRING (Szklarczyk et al., 2015). Genes involved in different biological pathways are colored differently. **(C)** RNA-Seq data for representative genes in *C. rota* (Schnytzer et al., 2018) and XBP1^*Flox*^ and XBP1^*LKO*^ mice. **(D)** RNA-Seq data for *Gabpa* in *C. rota*. Data are graphed as the mean ± SEM (n = 2).

## SUPPLEMENTAL TABLES

**Table S1. FPKM quantification of RNA-Seq data in XBP1^*Flox*^ and XBP1^*LKO*^ mice, related to Figure 1.** Ensemble gene ID, gene symbol, gene biotype and FPKM quantification of both duplicates and mean at each CT in XBP1^*Flox*^ and XBP1^*LKO*^ mice were shown.

**Table S2. Eigenvalue/pencil decomposition of all transcriptome with mean larger than 0.01 in XBP1^*Flox*^ and XBP1^*LKO*^ mice, related to Figure 1.** Period, decay rate, mathematical and biological phase and amplitude for each gene were provided as well as the mean value. Relative amplitude was calculated by normalizing the absolute amplitude to the mean.

Tab 1: Genes in XBP1^*Flox*^ mice

Tab 2: Genes in XBP1^*LKO*^ mice

**Table S3. Lists of genes with superimposed circadian rhythm in XBP1^*Flox*^ and XBP1^*LKO*^ mice, related Figure 1.** Circadian period, decay rate, mathematical and biological phase and amplitude for each gene were provided as well as the mean value. Relative amplitude was calculated by normalizing the absolute amplitude to the mean.

Tab 1: Genes in XBP1^*Flox*^ mice

Tab 2: Genes in XBP1^*LKO*^ mice

**Table S4. Lists of genes with superimposed ∼12h rhythm in XBP1^*Flox*^ and XBP1^*LKO*^ mice, related Figure 1.** ∼12h period, decay rate, mathematical and biological phase and amplitude for each gene were provided as well as the mean value. Relative amplitude was calculated by normalizing the absolute amplitude to the mean.

Tab 1: Genes in XBP1^*Flox*^ mice

Tab 2: Genes in XBP1^*LKO*^ mice

**Table S5. GO terms associated with the 4,600 genes whose 12h rhythms were either abolished or dampened in XBP1^*LKO*^ compared to XBP1^*Flox*^ mice, related to Figure 1.**

**Table S6. Lists of genes involved in RNA metabolism, protein metabolism, maintaining Golgi integrity and function, nucleotide metabolism and amino sugar/nucleotide sugar and N-glycan biosynthesis, whose 12h rhythms were either abolished or dampened in XBP1^*LKO*^ compared to XBP1^*Flox*^ mice, related to Figure 2.** ∼ 12h period, decay rate, mathematical and biological phase and amplitude for each gene were provided as well as the mean value. Relative amplitude was calculated by normalizing the absolute amplitude to the mean. If a ∼12h rhythm is not found, it is indicated by N/A.

Tab 1: Genes involved in RNA metabolism

Tab 2: Genes involved in protein metabolism

Tab 3: Genes involved in maintaining Golgi integrity and function

Tab 4: Genes involved in nucleotide metabolism

Tab 5: Genes involved in amino sugar/nucleotide sugar and N-glycan biosynthesis

**Table S7. Eigenvalue/pencil decomposition of all metabolites reported in (Krishnaiah et al., 2017), related to Figure 4.** Period, decay rate, mathematical and biological phase and amplitude for each metabolite were provided as well as the mean value. Relative amplitude was calculated by normalizing the absolute amplitude to the mean.

**Table S8. Eigenvalue/pencil decomposition of MMH-D3 transcriptome, related to Figure 5.** Period, decay rate, biological phase and amplitude for each gene were provided as well as the mean value. Relative amplitude was calculated by normalizing the absolute amplitude to the mean.

Tab 1: all genes

Tab 2: ∼12h genes

Tab 3: Circadian genes

**Table S9. GO terms associated with ∼12h-cycling genes in MMH-D3 cells, related to Figure 5.**

**Table S10. List of 12h-cycling CEDIF genes uniquely and commonly found in mouse liver *in vivo* and MMH-D3 cells *in vitro*, related to Figure 5.**

Tab 1: Commonly found ∼12h genes

Tab 2: Uniquely found ∼12h genes in vivo

Tab 3: Uniquely found ∼12h genes in vitro

**Table S11. Quantification and eigenvalue/pencil decomposition of XBP1s ChIP-Seq, related to Figure 6.** RPKM quantification and eigenvalue/pencil-identified 12h rhythm of XBP1s cistrome.

Tab 1: Promoter-associated XBP1s cistrome

Tab 2: Non promoter-associated XBP1s cistrome

Tab 3: List of genes bound by promoter-associated XBP1s with or without 12h transcriptome

**Table S12. List of genes with conserved 12h rhythms in both *C. rota* and mouse liver, related to Figure S7.** ∼12h period, decay rate, mathematical and biological phase and amplitude for each gene were provided as well as the mean value. Relative amplitude was calculated by normalizing the absolute amplitude to the mean.

**Table S13. GO terms associated with conserved 12h genes in mouse liver and *C. rota*, related to Figure S7.**

Tab 1: Using all mouse genes as background

Tab 2: Using all conserved genes (in mouse and *C. rota*) as background

